# Intraspecific variation and environmental filtering jointly and dynamically shaped individual elementomes of two sympatric small mammals

**DOI:** 10.64898/2026.08.31.748240

**Authors:** José R. Montiel-Mora, Vladislav Chrastný, Jiří Šindelář, Adéla Šípková, Markéta Zárybnická, Thibaut Rota

## Abstract

1. The biogeochemical niche hypothesis (BNH) recently proposed the multi-elemental composition (i.e., their elementome) as a promising ecological dimension. However, whether elementomes merely reflect species-specific stoichiometric constraints that vary little or randomly among individuals (the null BNH) or instead integrate structured ecological information (the alternative BNH), and, if the latter holds, how much elementome variation we can explain, remain unanswered questions. Our main hypothesis was that biological factors (species and intraspecific variation) would explain essential elementomes (Ca, Mg, Na, K, Zn, Cu, Mn, Fe, Mo, and Co), and that environmental filtering would explain non-essential elementomes (Pb, Al, V, Cr, Ni, As, Sr, Ba, Li, and B).
2. Here, we studied how species identity and intraspecific variation (body mass and sex), together with environmental filtering (season and habitat), contribute to shape the mandibular elementomes of eighty-four individuals belonging to two sympatric small mammals (*Apodemus flavicollis* and *Clethrionomys glareolus)*. We further evaluated ontogenetic body-mass allometric scaling in elemental stoichiometry under the vertebrate bone hypothesis (VBH).
3. We show that elementomes embed significant ecological information, as we explained up to 41.8% of multi-elemental variance. Species identity and intraspecific variation contributed more strongly to essential elementomes (19.7% of multi-elemental variance, vs. 11.8% for the environment). In contrast, environmental filtering contributed more strongly to non-essential elementomes (18.3% vs. 13.3% for intraspecific variation). Despite their phylogenetic proximity, ecological sympatry, and omnivorous diets, the species maintained distinct but moderately overlapping elementomes. Interestingly, the elementomes of the two species diverged from spring to autumn, perhaps as a result of micro-habitat or diet partitioning to reduce competition.
4. Ontogenetic body-mass scalings of several elemental and Ca-substitution relationships were consistent with predictions of the VBH. However, these scalings were context dependent, with, for instance, hypermetric scalings in autumn that could reflect life-history processes.
5. Our findings across individual and species levels of biological organization show that animal elementomes emerge from the combined influence of intraspecific variation, species identity, and environmental filtering, and that these factors explained a substantial proportion of multi-elemental variance. To conclude, elementomes provide useful ecological information on free-ranging animals.

## Introduction

The biogeochemical niche hypothesis (BNH) offers a framework for understanding how the elements on Earth and their human-driven disruption in the Anthropocene influence evolutionary and ecological processes, and how these processes, in turn, could shape biogeochemical niches (hereafter referred to as “elementomes”; Peñuelas et al., 2019). The BNH posits that an organism’s ecology (and by extension, community or ecosystem-level processes) could be approached through its elementome within a multi-elemental space (Peñuelas et al., 2019). Thence, elementomes have the premise to reflect the combined influence of metabolic and stoichiometric processes (e.g., resource acquisition, assimilation, storage, excretion) at intra- and interspecific levels of biological organisation, environmental exposure and bioavailability, and trophic and non-trophic interactions at community and ecosystem levels (Leroux, 2018; Peñuelas et al., 2019). In that respect, it is thought that elementome assembly is shaped not only by the geological context and elemental contamination but also by eco-evolutionary processes (González et al., 2018; Peñuelas et al., 2019).

The BNH has received increasing attention in plants (e.g., Liu et al., 2025; Sardans et al., 2021). However, the BNH (i.e., including a wider pool of elements than the classical C–N–P stoichiometric framework; Peñuelas et al., 2008, 2019) remains little tested in animals (but see Bartrons et al., 2018; Leroux, 2018; Moura et al., 2018; Němec et al., 2018; Zhang et al., 2022, 2025). The issue is especially relevant given the widespread metal(oid) contamination across ecosystems (Hanif et al., 2025). Mining, industrial emissions, agriculture, and fossil-fuel combustion continuously introduce potentially highly toxic and persistent non-essential metal(oid)s into the environment, contributing to their accumulation and potential transfer throughout food webs (Gall et al., 2015), thereby threatening to jeopardise ecosystems and human health (Hanif et al., 2025). Although this issue is well studied in environmental geochemistry and ecotoxicology (Al Sayegh Petkovšek et al., 2014; Martiniaková et al., 2010), it is rarely done through the ecological lens of the BNH (e.g., Bartrons et al., 2018; Zhang et al., 2022).

Intraspecific variation is a major driver of elemental stoichiometry (ES) (Nessel et al., 2024; Rota et al., 2026; Zhang et al., 2025). Ontogenetic variation in body mass induces changes in ES due to shifts in diet (Kraemer et al., 2012), metabolism (Elser et al., 2000), growth, ageing (Allen & Gillooly, 2009), skeletal development, and tissue maintenance (May & El-Sabaawi, 2024). Sex-specific physiology and reproductive investment may influence elemental accumulation and regulation differently between males and females throughout their ontogeny (Burger, 2007). Hence, seasonal changes in population size structure, reproduction, and environmental exposure, driven by natural and human activities, may interact with ontogeny in shaping species elementomes (Allison et al., 2025). For instance, increased fossil-fuel and coal combustion for winter heating, combined with frequent low clouds and fog, can enhance the deposition of airborne anthropogenic elements (Bridges et al., 2002). Non-essential elements could then accumulate in snowpacks and be released during late-winter snowmelt (Avak et al., 2019; Cimova et al., 2016), increasing their bioavailability during the breeding season. Therefore, distinguishing between essential and non-essential elements (Table S1) could be key to disentangle the relative contributions of intraspecific variation vs. environmental factors to elementome assembly (Jasiulionis et al., 2018). One would expect that the assembly of essential elementomes, because essential elements sustain biological processes (Table S1), is shaped primarily by evolutionary and ecological processes (e.g., intraspecific variability; El-Sabaawi et al., 2016; Rota et al., 2026), whereas the assembly of non-essential elementomes (composed of elements without known biological function; Table S1) would rather be the result of environmental filtering conditioning on the exposure to elements whose distributions are human-driven (Camizuli et al., 2018). Nevertheless, the ontogenetic ES of both essential and non-essential elements may be codependent and thus covary, for instance, via chemical substitutions for Calcium in bones (May & El-Sabaawi, 2024).

Bones are metabolically active, calcified tissues composed of 50–70% hydroxyapatite (Ca_10_(PO_4_)_6_(OH)_2_), and are fundamental to the structural, locomotive, and physiological functioning of vertebrates (Ciosek et al., 2021; May & El-Sabaawi, 2024). Skeletal tissues account for 80–90% of total mineral ash (Ciosek et al., 2021), and unlike visceral or muscular tissues, bones integrate ES over extended spatiotemporal scales (May & El-Sabaawi, 2024). To address key gaps in vertebrates’ ES, May & El-Sabaawi (2024) proposed the vertebrate bone hypothesis (VBH), suggesting that bones (1) drive whole-body vertebrate stoichiometry; (2) act as metabolically dynamic, flexible organs that regulate and store elements according to physiological state (i.e., breaking homeostasis’ assumption); and that (3) bone stoichiometry fluctuates across ontogeny, life history, and environmental contexts. Thus, a key yet little-tested general prediction of the VBH is the elemental flexibility of bones. Even at constant mineral content, bone stoichiometry fluctuates because foreign cations substitute for Ca^2+^ or PO_4_^3-^ in hydroxyapatite during crystal nucleation and turnover, particularly when Ca and P are scarce or environmental mimics abundant (Cazalbou et al., 2004; Glimcher, 2006; May & El-Sabaawi, 2024). Such substitutions are likely critical during ontogeny and reproduction, especially in metal-contaminated environments (Rodríguez-Estival et al., 2013). Because bone turnover and elemental accumulation are dynamic through ontogeny, we suggest here that elemental substitutions can manifest as power-law scalings relative to ontogenetic body mass. Therefore, we build upon the VBH to derive allometric predictions for elemental substitutions (*X_ij_/Ca*) across ontogeny, sex, and seasonal reproductive stages (Appendix 1).

Here, we studied elementome assembly under the BNH and VBH in two sympatric small mammal species, *Apodemus flavicollis* and *Clethrionomys glareolus*, using mandibles sampled from individual animals. The Pan-European distribution (Deffontaine et al., 2005; Michaux et al., 2004), high abundance, short life cycle, and intermediate position within food webs of these two omnivores make them promising vertebrate models to test both BNH and VBH frameworks, in addition to their use as bioindicators in metal-contaminated sites (Al Sayegh Petkovšek et al., 2014). These two species are exposed to toxic and non-toxic elements through multiple pathways, including ingestion of soil particles and uptake via their diet, which consists of seeds, plants, and invertebrates (Gall et al., 2015). Further, habitat differences may act as environmental filters: spruce-dominated forests are associated with more acidic soils than beech-dominated forests, potentially increasing the bioavailability of metals (Koptsik et al., 2023).

Our main goal was to assess how biological factors at inter- and intraspecific levels (species identity, ontogenetic variation in body mass, and sex) and environmental filters (season and habitat type) shape the assembly of essential and non-essential elementomes in two sympatric small mammal species. As a null hypothesis for the BNH, we would expect that elementomes are merely reflecting species-specific stoichiometric constraints imposed by their structural morphology and/or evolutionary history (e.g., 70-90% of leaf elementome variation in trees is driven by phylogeny; Sardans et al. 2021). In other words, this null hypothesis suggests that elementomes vary little among individuals, or vary in an unstructured, random way among individuals within species or among ecological contexts spanned by those individuals (ontogenetic stages, sex, habitats, seasons, etc.). Since trait-based ecology is built on the assumption that traits are functional (i.e., they relate to fitness and ecological dynamics, and can be measured at the individual level; Rota et al. 2026 and references therein), accepting the null BNH would limit the integration of the elementome to existing trait-based frameworks, relaying its use to macroecological studies at the interspecific level (e.g., Sardans et al. 2021). Alternatively, we expected that elementomes, in addition to reflecting species-specific stoichiometric constraints, would embed important ecological information at the biological level of individuals, thus showing promise for integrating elementomes as a novel trait-based source of information in ecology.

Specifically, we tested four hypotheses. First, (H1), we expected the relative contributions of biological and environmental factors to differ between essential and non-essential elementomes. Because essential elements sustain biological processes and are expected to be under stronger physiological regulation, we predicted that biological factors (species identity, body mass, and sex) would contribute more strongly to essential elementomes, whereas environmental filters (season and habitat type) would contribute more strongly to non-essential elementomes. Second (H2), we hypothesised that elementomes differ between species, reflecting evolutionary, ecological, and physiological differences. Given that elementome assembly likely encompasses a large array of biological and environmental processes, we expected that full elementomes would embed high dimensionality (Zhang et al., 2022). However, while we expected significant niche differentiation between species, their sympatry, phylogenetic proximity, and broadly similar omnivorous diets led us to expect some overlap in their elementomes. Third (H3), we hypothesised that ontogenetic variation in body mass, in addition to life-history related to reproductive constraints (sex and season), contributes to shape elementomes, with stronger associations for essential elementomes. Within this hypothesis, we tested specific predictions of ontogenetic scaling under the VBH (May & El-Sabaawi, 2024; see Appendix 1 and predictions *p1*-*p4*). Fourth, (H4) we expected environmental variation to shape mandibular elementomes. Specifically, we predicted higher burdens of non-essential elements in spring than in autumn because metal(oid)s accumulated in snowpacks may be released during snowmelt, as well as shifts in elementomes between spruce and beech forests associated with differences in soil chemistry and metal(oid)s contamination. Finally, we tested for seasonal changes in elementome partitioning and discussed implications for the coexistence of these two sympatric species.

## Materials and methods

### Study sites

We conducted field sampling in spring (early June) and autumn (October) 2017, in the Ore Mountains (50°41′ N, 13°36′ E, elevation: 735-960 m a.s.l., Czechia), within the European Black Triangle. This region experienced substantial atmospheric emissions from brown coal (lignite) combustion and industrial activities in the 1960s–1990s (Marx & McGowan, 2010), resulting in extensive metal contamination and forest diebacks (Hošek et al., 2024). Soils in our location show high concentrations of potentially toxic elements, including As and Pb (Figure S1; Kochergina et al., 2017). We sampled small mammals in two habitat types: spruce (*Picea* spp.) and beech (*Fagus sylvatica*) dominated forests surrounded by open grassland habitats, which differ in element exposure (Figure S1) and pH (spruce forest: 3.75 ± 0.06 sd; beech forest: 4.33 ± 0.12 sd).

### Small mammal trapping design

We collected small-mammal samples within five trapping plots, each covering 0.27 ha, and conducted trapping for three consecutive days during each sampling event. Within each plot, we arranged 40 snap traps in a regular 4 × 10 grid, with ∼10 m between traps. We baited traps with a mixture of flour, fat, and roasted bacon (Zárybnická et al., 2017). We identified captured individuals into species and measured their body mass using a Pesola spring scale. We determined sex by examining the reproductive organs, identifying males by their testes and females by their uterus. We kept specimens on ice packs in cool boxes during transport to the laboratory, where we stored them frozen until laboratory processing. We conducted the research in accordance with national and international ethical standards and Czech national legislation. Small mammal trapping was conducted as part of authorised wildlife monitoring activities in the study region under a permit issued by the Ministry of the Environment of the Czech Republic (No. 71735/ENV/16-3580/630/16).

### Metal and metalloid analysis

We analysed the left and right mandibles from each individual as a single sample, after removing teeth and surrounding soft tissues. We digested the samples and quantified elemental concentrations by ICP-OES and ICP-MS; full sample-preparation, instrumental, calibration, and quality-control procedures are provided in Table S2. We quantified ten essential elements (Ca, Mg, Na, K, Zn, Cu, Mn, Fe, Mo, and Co) and ten non-essential elements (Pb, Al, V, Cr, Ni, As, Sr, Ba, Li, and B), expressed as mg kg⁻¹ (ppm), whose rationale for classification as essential and non-essential elements is presented in Table S1.

### Statistical analysis

After removing two individuals lacking sex and body mass data, our study comprised 84 individuals: 45 *A. flavicollis* and 39 *C. glareolus*. Sample sizes by species, season, sex, and habitat type are provided in Table S3. We performed all statistical analyses and graphics in R 4.5.1 (R Core Team, 2025).

#### a. Multi-elemental analyses

Multivariate analyses were conducted using the “vegan” R package (Oksanen et al., 2001). We treated essential and non-essential elements as separate datasets. After confirming the absence of zero or negative concentrations, we applied a centred log-ratio (*clr-*) transformation and calculated Euclidean distances on *clr-*transformed data to obtain an Aitchison distance matrix, used for compositional analysis (i.e., avoiding the bounded bias inherent to concentration data; Aitchison, 1982). Hence, we used this distance for PERMANOVA and PERMDISP analyses.

We first conducted separate principal component analyses (PCAs) on *clr*-transformed essential and non-essential compositions. For all PCAs, we retained as many PCs as needed to reach ca. 70% of variance explained. Body mass was ordinated as a supplementary variable onto the first three PCA axes using *envfit* with 999 permutations for graphical representations of PCAs. For each species separately, we tested log_10_-transformed body mass, sex, season, and habitat, and their two-way interactions using sequential PERMANOVA (*adonis2*; 999 permutations). Pooled-species PERMANOVAs included species, species-specific effects of body mass, season, sex, and habitat type, and the specified three-way interactions, whereas species-specific models excluded species. Homogeneity of multivariate dispersion for categorical predictors was assessed using the *betadisper* and *permutest* functions.

To evaluate H1, we summarised sequential PERMANOVA R² contributions into biological, environmental, interaction, and residual components. In pooled analyses, the biological component comprised species, body mass × species, and sex × species, whereas the environmental component comprised season × species and habitat type × species; the remaining interaction terms formed the interaction component. In species-specific analyses, biological predictors were body mass and sex, whereas season and habitat type were environmental predictors.

To evaluate H2, we additionally analysed the complete elementome, expecting high dimensionality (Zhang et al., 2022), which we quantified by the number of PCs retained. We quantified species’ distance to centroid and within-species dispersion in this retained space and calculated species-specific convex-hull overlap in the PC1–PC2 projection. We then constructed a bi-dimensional space based on log10-transformed Ba/Ca and Sr/Ca ratios and calculated the same metrics. The Ba/Ca vs. Sr/Ca space has been established as a proxy of trophic niches in terrestrial ecosystems, from plants to herbivores and predators (Balter, 2004). Thus, comparing the discriminatory power of the elementome and Ba/Ca vs. Sr/Ca spaces constituted a rough test of the BNH. Species niche differences and dispersion were tested using PERMANOVA and PERMDISP, respectively, with 999 permutations. Potential association between the two distance matrices was assessed using a Pearson Mantel test with 999 permutations.

To evaluate H4, we examined the effects of season and habitat in the multivariate models. To further examine seasonal species elementome partitioning, we conducted follow-up PERMANOVAs comparing species within each season separately for essential and non-essential elementomes, and quantified interspecific overlap between 95% PCA ellipses as the intersection area relative to their union.

#### b. Ontogenetic scaling of elemental stoichiometry

To evaluate H3, we first considered the total pool of elements and conducted species- and element-specific allometric linearised models (using ordinary least squares ‘OLS’ regressions) relating log10-transformed elemental concentrations to log10-transformed body mass while accounting for season and habitat type. These models were also used to evaluate element-specific season and habitat effects under H4. Then, to evaluate the VBH predictions suggested in Appendix 1, we fitted allometric linearised models using robust *rlm* regression (to reduce the bias inherent to low sample sizes) on a restricted pool of divalent cations (Pb²⁺, Sr²⁺, Ba²⁺, Mg²⁺, and Ca²⁺) and their ratios with Ca, to evaluate elemental substitutions for Ca (Pb/Ca, Sr/Ca, Ba/Ca, Mg/Ca). For each species, one model set allowed body-mass scalings to vary between sexes, and the other allowed them to vary between seasons.

## Results

*A. flavicollis* had a higher mean body mass than *C. glareolus* (mean ± SD: 27.4 ± 7.07 g; range: 14–44 g; n = 45 versus 19.2 ± 4.11 g; range: 13–33 g; n = 39, respectively). Calcium dominated mandibular composition in both species, with 26.5% in *A. flavicollis* and 27.3% in *C. glareolus*, followed by Mg (0.24%; 0.23%) and Na (0.24%; 0.21%), whereas Ba (0.036%; 0.032%), Zn (0.031%; 0.034%), and Sr (0.034%; 0.024%) showed minor contributions (Table S4). *C. glareolus* had higher mean concentrations for the non-essential elements Al, Pb, and V, and for the essential elements Fe, Mn, and Zn. In contrast, *A. flavicollis* had higher mean concentrations for the essential elements Mg and Na, and for the non-essential elements Sr and Ba (Table S4).

### Biological and environmental drivers of elementome assembly

According to our first hypothesis (H1), we observed a dichotomy between the relative contributions of biological and environmental factors to essential and non-essential elementome assembly (Figure 1; variance explained ranged from 30–41.8%). When considering both species, biological factors explained more variation than environmental factors in essential elementomes (19.7% vs. 11.8%, respectively), whereas environmental factors explained more variation in non-essential elementomes (18.3% vs. 13.3%; Figure 1; Table S5). In *A. flavicollis*, environmental factors were markedly stronger predictors of non-essential elementomes than biological factors (21.6% vs. 8.0%), whereas in *C. glareolus* environmental factors explained more variation in both elemental groups (Figure 1; Table S5).

**Figure 1.**
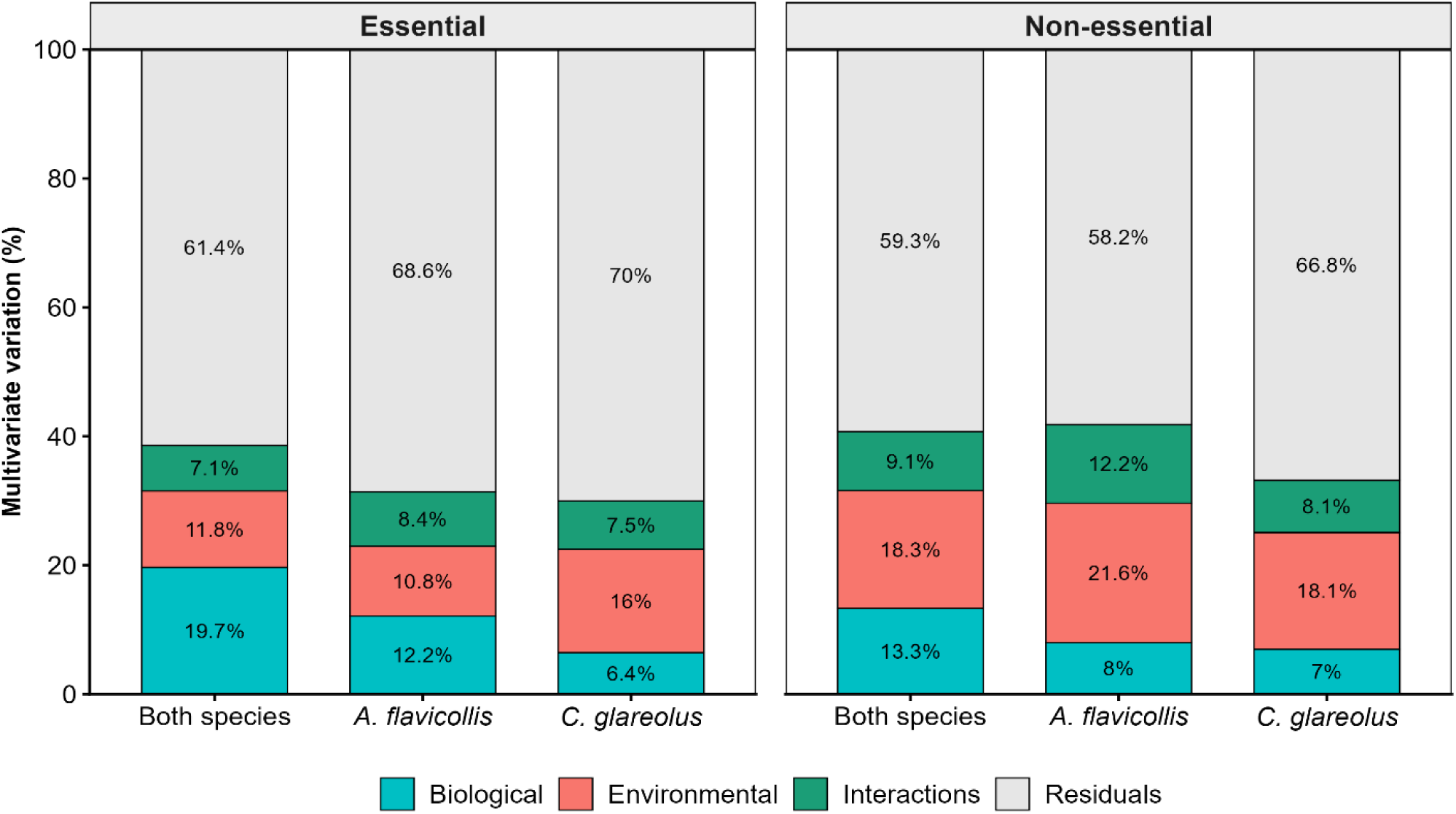
Relative contributions of biological, environmental, interaction, and residual components to variation in essential and non-essential mandibular elementomes. Percentages represent R² contributions from sequential PERMANOVA models for both species combined and for *A. flavicollis* and *C. glareolus* separately.

### Interspecific variation in the mandibular elementome

Consistent with our second hypothesis (H2), *A. flavicollis* and *C. glareolus* showed a moderately overlapping elementome, with 95% PCA ellipse overlap of 37.5% for essential and 58.7% for non-essential elements (Figure 2), with significant species differences for both elemental groups (Table S6). Multivariate dispersion differed between species for the non-essential elementome, with *C. glareolus* individuals showing a greater mean distance to the centroid than *A. flavicollis* individuals (1.30 vs 1.05, respectively; PERMDISP *p* = 0.023). The first two PCs explained 63.9% and 50.4% of the variation in essential and non-essential elementomes, respectively (Figure 2a, b), and the explained variance of essential and non-essential elementomes increased to 85.2% and 70.1% when the first three PCs were considered (Figure S2).

**Figure 2.**
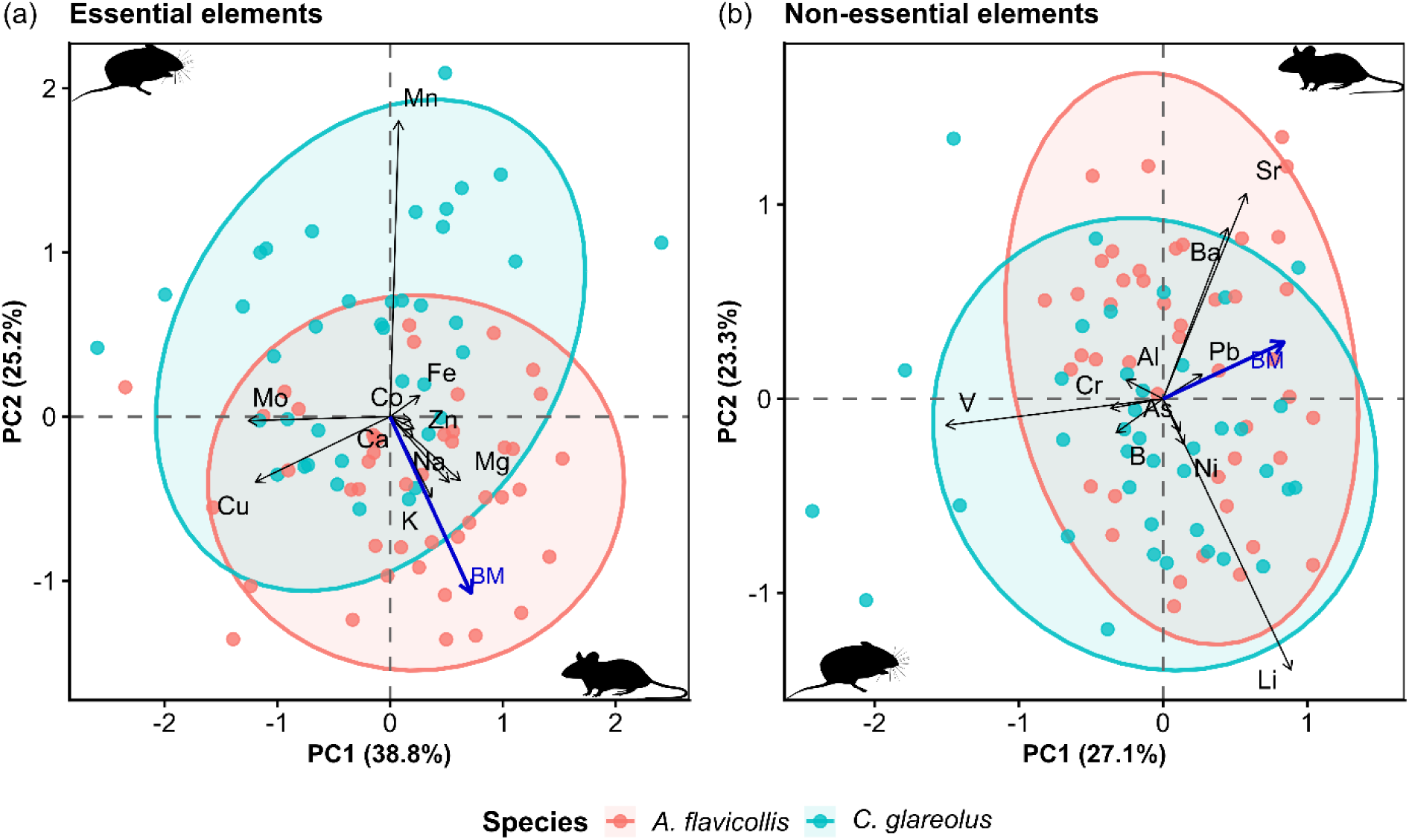
PCAs of *clr*-transformed elemental compositions for (a) essential and (b) non-essential elements. Points represent individuals, colours indicate species, and ellipses represent 95% ellipses for each species. Black arrows show element loadings, and the blue vector shows the significant association of body mass with the ordination, fitted as a supplementary variable (not contributing to the ordination). Animal silhouettes were obtained from ‘PhyloPic.org’ (public domain).

When considering the full elementome (all 20 elements), the first five PCs explained 74.8% of the total variation, consistent with our expectation of high elementome dimensionality (i.e., here at least five orthogonal dimensions; Zhang et al., 2022). Both the retained multi-elemental space and the Ba/Ca vs. Sr/Ca bi-dimensional space showed significant species differentiation, with no evidence of unequal multivariate dispersion (Table S7; Figure S3). The multi-elemental representation showed a slightly higher PERMANOVA R² and lower convex-hull overlap than the ratio-based space (Table S7; Figure S3). The two distance matrices were weakly but significantly correlated (Mantel *r* = 0.231; *p* = 0.003), indicating a partial association between the two elemental distances. Overall, these findings supported the BNH (H2) by suggesting that elementomes capture fine-scale interspecific differentiation while embedding high ecological dimensionality (Zhang et al., 2022).

### Intraspecific variation in the mandibular elementome

Consistent with our third hypothesis (H3), body mass affected the elementome of both species (Figure 2; Table S6). Species-specific models showed significant body-mass effects on both element groups in *A. flavicollis*, whereas in *C. glareolus*, body mass significantly explained variation in the non-essential elementome but not the essential elementome (Figure 2; Table S6). In contrast, sex-related effects were not significant in either element group in the global or species-specific PERMANOVAs (Table S6).

Body mass allometric scalings for individual elements were more often significant in *A. flavicollis* than in *C. glareolus* (Figure 3; Table S8). In *A. flavicollis*, Mg and Na showed positive allometric scalings (*b* = 0.716 and 0.712, respectively), whereas Ca, Cu, Mo, Al, Cr, and V showed negative allometric scalings (*b* ranging from –0.155 to –1.022; Table S8). In *C. glareolus*, Mg and Na scaled allometrically-positive with body mass (*b* = 0.731 and 0.46, respectively), whereas Mn scaled hypermetrically-negative (*b* = –1.312; Table S8).

**Figure 3.**
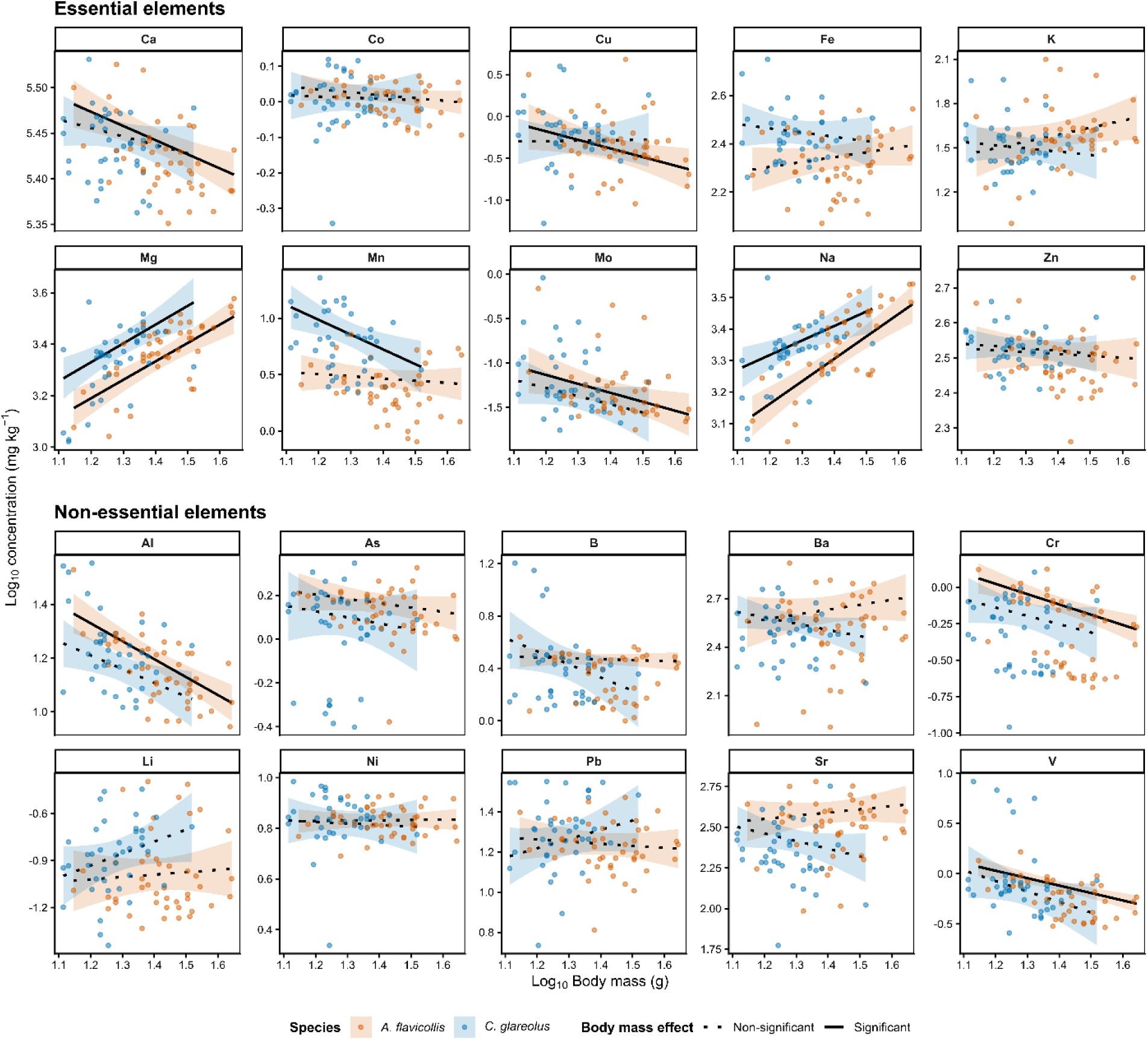
Log-log allometric relationships between body mass and mandibular elemental concentrations in *A. flavicollis* (orange) and *C. glareolus* (blue). Points represent individual observations. Solid black lines indicate significant relationships, while dotted black lines indicate non-significant relationships. Shaded bands are 95% confidence intervals.

The robust allometric scalings were broadly consistent with the predictions derived from the VBH and outlined in Appendix 1 (Figure S4; Figure S5, Tables S9-S10). Consistent with *p1*, Mg concentrations scaled from positively-allometric to positively-hypermetric with body mass for both sexes and species (*b* ranging between 0.644 and 1.268; Figure S5; Table S9). Pb, Sr, and Ba showed shallower but generally positive scalings in females (Figure S5; Table S9). Consistent with *p2*, Ca tended to scale negatively with body mass, especially in *A. flavicollis* (Figure S5; Table S9), although we did not detect strong scaling differences between sexes. Supporting *p3*, X_ij_/Ca ratios scaled positively with body mass in females, whereas Sr/Ca and Ba/Ca showed negative relationships in males (Figure S4; Table S10). Contrary to *p4*, several hypermetric scalings occurred in autumn (*b* up to 2.061; Figure S4; Table S10), although this pattern varied among elements and species. Overall, these results supported H3 and the VBH by showing that mandibular elemental stoichiometry is highly dynamical across body mass, sex, and seasonal contexts.

### Environmental filtering of elementome assembly

#### a. Effects of seasons

Consistent with our fourth hypothesis (H4), seasons significantly shaped both essential and non-essential elementomes, with a slightly higher association with non-essential elementomes (Table S6). According to our prediction, spring was associated with higher concentrations of the essential elements Ca, Co, and Mn and the non-essential elements Al, As, B, Ba, Cr, Li, Ni, Sr, and V in *A. flavicollis* (Figure S6; Table S11). In *C. glareolus*, spring was associated with higher concentrations of Ca, As, and Cr (Figure S6; Table S11).

Species-specific PERMANOVAs showed significant seasonal effects on both element groups (Table S6). Post hoc analyses showed that elementome partitioning between species in autumn was significant for both essential (PERMANOVA, F = 7.64, R² = 0.193, *p* = 0.001) and non-essential elementomes (F = 4.48, R² = 0.123, *p* = 0.001). In spring, partitioning was weaker than in autumn but remained significant for the essential elementome (F = 5.89, R² = 0.109, *p* = 0.001), while it was not significant for the non-essential elementome (F = 2.24, R² = 0.045, *p* = 0.057; Figure S7). Consistently, interspecific overlap was lower in autumn than in spring for essential (23.3% vs. 37.1%; Figure S7) and non-essential elementomes (33.5% vs. 51.8%; Figure S7). Overall, elementome partitioning between the two sympatric species increased from spring to autumn for both essential and non-essential elementomes.

#### b. Effects of habitats

Habitat type significantly shaped non-essential elementomes in both species, whereas significant effects on the essential elementome were detected only in *C. glareolus* (Table S6). Contrary to our fourth hypothesis (H4), habitat responses were limited in *A. flavicollis*, with only Fe showing higher concentrations in beech forests (Figure S8; Table S11). In contrast, *C. glareolus* showed significant habitat responses for Mg, Na, Ba, and Sr that exhibited higher burdens in beech forests, whereas Al and V showed higher burdens in spruce forests (Figure S8; Table S11). Therefore, the magnitude and direction of these responses were contingent on species and elements, with *C. glareolus* showing greater responsiveness to habitat types than *A. flavicollis* (Figure S8; Table S11).

## Discussion

We show here that mandibular elementome assembly in two sympatric small mammals is shaped by the combined influence of intraspecific variation and environmental filtering, with their relative contributions differing between essential and non-essential elementomes. Biological factors contributed more strongly to essential elementome assembly, whereas environmental factors contributed more strongly to non-essential elementomes. Species identity contributed to interspecific elementome differentiation, consistent with the BNH (Peñuelas et al., 2019), whereas ontogenetic variation in body mass emerged as a strong driver of mandibular ES among individuals, consistent with the VBH (May & El-Sabaawi, 2024). Overall, our findings support the BNH as an integrative framework for interpreting animal elementomes and highlight the relevance of the VBH for understanding bone ES dynamics in wild animals, while emphasising the importance of intraspecific variation to shape elementomes (Nessel et al., 2024).

### Biological and environmental drivers of elementome assembly

Our partitioning analysis suggests that elementome assembly emerged from the combined influence of physiological regulation, environmental exposure, and their interaction, rather than from either process alone. Across both species, biological factors (species identity and intraspecific variation) explained a greater proportion of variance in essential elementomes, whereas environmental factors (season and habitat type) rather shaped non-essential elementomes (Moura et al., 2018). However, the effects of biological and environmental factors we observed were species-dependent, with environmental factors contributing more strongly to elementome assembly in *C. glareolus* than in *A. flavicollis*, suggesting that the former may be more useful as a bioindicator of elemental exposure (Al Sayegh Petkovšek et al., 2014). Nevertheless, most of the elementome variation remained unexplained, likely because elementomes integrate processes not captured here (Peñuelas et al., 2019), highlighting the need for further integrative research combining trait-based approaches (Rota et al., 2026). Understanding the high flexibility of bone elementomes will require the integration of elemental dynamics in bones alongside biological traits, life histories, and environmental contexts (Jota-Baptista et al., 2022; Němec et al., 2018).

### Interspecific differentiation in mandibular elementomes

Confirming our second hypothesis, *A. flavicollis* and *C. glareolus* occupied distinct positions in both multi-elemental and Ba/Ca–Sr/Ca spaces despite their phylogenetic proximity, sympatry, and broadly similar omnivorous diets (Abt & Bock, 1998). Importantly, this indicates that even closely related, ecologically similar, sympatric mammals can maintain distinct elementomes. This differentiation suggests that elementomes can capture relatively subtle ecological and physiological differences between closely related species, consistently with the BNH (Peñuelas et al., 2019). At the same time, the moderate overlap between species indicates that species identity does not impose a rigid elemental signature, leaving substantial scope for individual-level variation within species, consistent with growing evidence that intraspecific variation among individuals can be an important component of an animal’s ecology (Rota et al., 2026; Zhang et al., 2025). The high dimensionality of the complete elementome further suggests that elementome variation cannot be reduced to a small number of elemental axes (Zhang et al., 2022). This is particularly relevant given the ecological similarity of the two species, as it implies that multiple, orthogonal elemental dimensions contribute to their differentiation. Such multidimensionality may arise because different elements integrate distinct physiological functions, exposure pathways, and environmental constraints, allowing the elementome to capture ecological information that would be overlooked by individual elements or single elemental ratios.

Mandibular ES also differed between species in several trace elements and macroelements, consistent with species-specific elemental profiles previously reported in European small mammals inhabiting contaminated environments (Al Sayegh Petkovšek et al., 2014; Martiniaková et al., 2015). The partial concordance between the multi-elemental space and the Ba/Ca–Sr/Ca bi-dimensional space (Table S7; Figure S3) suggests that some of the interspecific structure captured by the elementome may overlap with variation represented by these trophic niche proxies (Balter, 2004). The niche overlap observed in both representations also indicates substantial individual-level variation within species, which may contribute to the similarity between species in the multi-elemental space. Interestingly, despite their similar ecologies and sympatry, the two species showed substantial elementome partitioning that increased from spring to autumn. Elementome partitioning in autumn, when abundances of small mammals are often higher, may reduce the competition of the behaviourally dominant *A. flavicollis* on the subordinate *C. glareolus* (Kurek et al., 2026), and thus prevent the competitive exclusion of the latter (Benedek & Sîrbu, 2025). Such seasonal partitioning of elementomes may therefore reflect ecological processes contributing to species coexistence. Overall, these findings are consistent with the BNH, whereby species-specific ecological strategies, physiological constraints, and environmental exposure jointly contribute to shape dynamically elementomes (Peñuelas et al., 2019), even for phylogenetically close, sympatric species sharing an omnivorous diet (Abt & Bock, 1998).

### Body mass allometric scalings of mandibular elemental stoichiometry

Body mass emerged as an important source of intraspecific variation in mandibular elementomes. In both species, Mg and Na increased with body mass, whereas Ca and several other elements declined, particularly in *A. flavicollis*. These contrasting patterns are consistent with variation in bone growth, mineral balance, tissue remodelling (Robling et al., 2006), and growth dilution (Zhang et al., 2025), whereby bone growth may outpace the incorporation or retention of particular elements, or conversely, elemental incorporation may increase faster than bone growth (May & El-Sabaawi, 2024; Zhang et al., 2025). Such allometric responses are consistent with the dynamic role of bone proposed by the VBH (May & El-Sabaawi, 2024). However, body mass should not be interpreted here as a direct measure of ontogenetic age (Peters, 2012; Viro & Sulkava, 1985). Instead, the body-mass relationships observed here are best interpreted as evidence of size-associated bone elemental flexibility.

The robust allometric analyses were broadly consistent with several predictions in Appendix 1, although support varied across elements, species, sexes, and seasons. Mg concentrations and Mg/Ca ratios generally increased from allometric to hypermetric with body mass, consistent with predictions *p1* and *p3*, whereas the decline in Ca observed in *A. flavicollis* was consistent with *p2*. Such contrasting changes in Ca and substitution cations may amplify X_ij_/Ca scaling because these ratios integrate variation in both the substituting cation and the Ca pool (May & El-Sabaawi, 2024; Appendix 1). Sr/Ca and Ba/Ca relationships varied among species, sexes, and seasons, providing partial support for the expectation of stronger elemental substitution patterns in females (Ciosek et al., 2021). Such sex-dependent patterns may be related to reproductive mobilisation of bone Ca, which can alter mineral balance and potentially modify the relative incorporation or retention of foreign cations in bone (Ciosek et al., 2021; May & El-Sabaawi, 2024). However, contrary to *p4,* several hypermetric scalings occurred in autumn rather than in spring. These strong departures from mass-invariant scaling suggest that bone elemental composition can change disproportionately across body mass and may reflect delayed effects of reproductive investment, seasonal changes in elemental exposure, or variation in bone remodelling and mineral incorporation (Robling et al., 2006; May & El-Sabaawi, 2024).

More generally, the influence of body mass on elemental composition depended on seasonal context, but significant seasonal interactions were restricted to Ba and Sr in *A. flavicollis* and Mg, Na, and Al in *C. glareolus*. Because both the direction and magnitude of these relationships varied among elements and species, our results do not support a general seasonal tendency for larger individuals to consistently accumulate higher or lower elemental concentrations. Instead, these species-, element-, and season-dependent responses reinforce the view that bone elementome assembly is context dependent and cannot be explained by body mass alone. These context-dependent responses of intraspecific stoichiometric scalings could shape the covariations among elements within species, leading to the emerging apparent interspecific patterns in elemental composition (El-Sabaawi et al., 2014), a research idea that will need further attention.

### Environmental variation in mandibular elementomes

We found that the environmental factors (season and habitat) were more strongly associated with non-essential than essential elementomes. This contrast is consistent with tighter physiological regulation of essential elements and greater environmental responsiveness of non-essential elements (Camizuli et al., 2018). Seasonal effects were broader in *A. flavicollis*, with spring associated with higher concentrations of several essential and non-essential elements, whereas *C. glareolus* showed fewer element-specific responses. The predominance of higher concentrations in spring, particularly for non-essential elements, is broadly consistent with the seasonal release of atmospherically deposited metals accumulated in snowpacks during winter and mobilised during snowmelt (Avak et al., 2019; Cimova et al., 2016). Another possibility would be that spring elemental compositions reflect the high growth rates in spring and bone mineral incorporation (Viro & Sulkava, 1985).

Habitat affected the elementome of both species, with stronger effects in *C. glareolus* than in *A. flavicollis*. Differences in pH and atmospheric deposition loads between habitats (Figure S1) observed in our study may have influenced elemental bioavailability and exposure (Bing et al., 2016; Koptsik et al., 2023). Here, we interpreted environmental filtering as the context-dependent influence of environmental conditions on elementomes, while recognising that these effects may also interact with species-specific ecological and biological characteristics. This interpretation is consistent with broader community-assembly frameworks in which environmental filtering operates alongside biotic processes rather than independently of them (Kraft et al., 2015). Incorporating such biotic interactions more explicitly represents an important next step forwards to extend the BNH to community ecology.

The mandibular elementome represents a time-integrated record (May & El-Sabaawi, 2024) that can be recovered from museum collections (DuBay et al., 2025) or natural remains (e.g., from bird of preys’ rejection pellets or paleological records; Scott et al., 2024), whose temporal integration may depend on species-specific metabolic rates (Allen & Gillooly, 2009). The high metabolic rate of small mammals would imply a highly dynamical mandibular bone stoichiometry, explaining the high flexibility we observed across ontogeny, seasons, and habitats in the individuals of two species of small mammals. Overall, our findings support the BNH as a general framework for understanding how ecological and environmental processes shape animal elementomes, and the VBH as a complementary perspective on how bone physiology contributes to variation in ES across individuals and species.

## Acknowledgements

We thank Nikola Gärtner for laboratory assistance and other bachelor’s and master’s students for their assistance with fieldwork. We gratefully acknowledge financial support from the Czech Science Foundation (GAČR) through research project No. 25-17362S.

## Author Contributions

**J. R. M-M.**: Performed statistical analyses, prepared the visualisations, and led the writing and revision of the manuscript. **T. R.**: Conceptualised the study, designed the questions and statistical analyses, and guided the manuscript’s review and editing. **M. Z.:** Initiated, conceptualised, and led the study, supervised specimen preparation for elemental analyses, and provided critical input throughout the development and revision of the manuscript. **V. C.** and **A. Š.**: Led and developed the analytical component of the study and performed all elemental analyses. **J. Š.**: Performed fieldwork and specimen collection. All authors edited the manuscript and approved the final submitted version.

## Data availability statement

The data and R code to reproduce the study are made publicly available in a Figshare repository (Montiel-Mora et al. 2026, DOI: 10.6084/m9.figshare.33340470; URL: https://doi.org/10.6084/m9.figshare.33340470).

