## Supplementary Material for "Intraspecific variation and environmental filtering jointly and dynamically shaped individual elementomes of two sympatric small mammals"

### *Appendix 1: Allometric Scaling of Ontogenetic Elemental Substitutions ($X_{ij}$/Ca)*

Over 90% of mammal's calcium (Ca) is stored as skeletal hydroxyapatite. Across ontogeny – especially during gestation and lactation in females – Ca is heavily mobilised from bones to maintain serum Ca homeostasis (Ciosek et al., 2021). Under the VBH (May & El-Sabaawi, 2024), we suggest how this decreasing Ca baseline could influence ontogenetic scaling of exogenous divalent substitution cations (e.g., $Sr^{2+}$, $Ba^{2+}$, $Mg^{2+}$, $Pb^{2+}$) directly substituting to $Ca^{2+}$ in bones’ hydroxyapatite.

We assume that the element intake in bones through time $I_i$ is scaling with body mass $M$ at ~ $M^{3/4}$, and that bone mass scales isometrically with $M_{bones}$ ~ $M^1$. At a steady state, when element accumulation in bones roughly equals element clearance rate ($I_i \sim k_i$), the concentration $i$ of any element scales as a power-law of body mass $M$:

$$[i] = a_i M^{bi}$$

A null slope ($b_i = 0$) means that element accumulation matches bone growth; concentrations are mass-invariant. A positive allometry ($b_i > 0$) means that the element accumulating in bones is outpacing bone’s growth (e.g., when $I_i > k_i$ and $k_i \sim 0$); the element concentration increases with body mass. A hypoallometry ($b_i < 0$) means that bone growth outpaces element accumulation; concentrations decrease with body mass. Drawing on substitution cations $X_{ij}$ chemistry and expected Ca decrease with ontogeny and reproduction, we predict (***p1*** to ***p2***):

***p1***: Divalent cations $X_{ij}$ ($Pb^{2+}$, $Sr^{2+}$, $Ba^{2+}$, $Mg^{2+}$) are replacing permanently $Ca^{2+}$ in the hydroxyapatite crystals through ontogeny, so that $I_i > k_i \sim 0$, leading to bioaccumulation ($b_{Xij} > 0$).

***p2***: Calcium ($Ca^{2+}$) is expected to have a higher balance towards clearance rate with age in females, so that $I_i < k_i > 0$ and $b_{Ca} < 0$.

Therefore, we suggest that when an exogenous cation $X_{ij}$ is substituting permanently for Ca, the elemental concentration ratio $[X_{ij}]$ / $[Ca]$ scales as a power-law:

$$\frac{[X_{ij}]}{[Ca]} = \frac{a_X M^{bx}}{a_{Ca} M^{b_{Ca}}} = \alpha_0 M^{b_{ratio}} \Leftrightarrow \ln\left(\frac{[X_{ij}]}{[Ca]}\right) = \ln(\alpha_0) + b_{ratio} \ln(M)$$

Because $b_{ratio} = b_X - b_{Ca}$, the physiological state of structural calcium directly modulates the slope of the foreign element ratio, such as if $b_{Ca} \sim 0$, $b_{ratio} = b_X - 0 = b_X$, and if $b_{Ca} < 0$, $b_{ratio} = b_X - (-|b_{Ca}|) = b_X + |b_{Ca}|$. We thus get the following predictions (***p3*** to ***p4***):

***p3***: For substitution cations $X_{ij}$, $b_{ratio}$ scales allometrically with body mass (>> 0 and/or > 1).

***p4***: For $X_{ij}$, $b_{ratio}$ (>> 0 and/or > 1) of female in spring during gestation/lactation > female in autumn > males (where $b_{ratio} \sim b_X$).

---

**Table S1.** Classification of analysed elements into essential and non-essential categories based on their biological roles and potential toxic effects in vertebrates.

| Category | Element | Biological function | Rationale | References |
|---|---|---|---|---|
| Essential (macroelements) | Ca | Bone mineralisation and structural integrity | Major component of hydroxyapatite in mineralised tissues | (Suttle, 2022) |
| | Mg | Enzyme cofactor and bone metabolism | Required for enzymatic activity and bone remodelling | (Rude et al., 2003) |
| | Na | Osmoregulation and nerve function | Maintains extracellular fluid balance and electrochemical gradients | (Michell, 1989) |
| | K | Cellular function and acid–base balance | Regulates membrane potential and acid–base balance | (Pohl et al., 2013) |
| Essential (trace elements) | Fe | Oxygen transport and cellular respiration | Component of haemoglobin and redox enzymes | (Ganz & Nemeth, 2006) |
| | Zn | Enzyme activity and protein structure | Cofactor in numerous enzymes involved in DNA synthesis and bone metabolism | (Cai et al., 2025) |
| | Cu | Redox reactions and connective tissue formation | Cofactor in redox enzymes and iron metabolism | (Lutsenko et al., 2025) |
| | Mn | Enzyme activity and antioxidant defence | Involved in bone formation and superoxide dismutase activity | (Taskozhina et al., 2024) |
| | Mo | Enzyme cofactor | Cofactor for oxidase enzymes involved in metabolic processes | (Novotny & Peterson, 2018) |
| | Co | Component of vitamin B12 | Essential constituent of cobalamin (vitamin B12) | (Banerjee & Ragsdale, 2003) |
| Non-essential (toxic) | Pb | None | Accumulates in bone and is associated with toxic effects | (Hydeskov et al., 2024) |
| | As | None | Toxic metalloid that interferes with cellular processes | (Ismail & Roberts, 1992) |
| Non-essential (analogues) | Sr | None | Calcium analog incorporated into the bone matrix | (Panahifar et al., 2019) |
| | Ba | None | Calcium analog with limited biological retention | (Panahifar et al., 2019) |
| Non-essential (other) | Al | None | Associated with environmental exposure and potential toxic effects | (D'Haese, 2013) |
| | Li | None | May influence cellular signalling | (Hart, 2024) |
| | B | None | May affect mineral metabolism | (Blagojevic & Pavlovic Ivan, 2025) |
| | V | None | May interfere with phosphate metabolism | (Etcheverry et al., 2012) |
| | Ni | None | May interfere with metal homeostasis | (Nielsen, 2021) |
| | Cr | None | Reported metabolic effects remain controversial in mammals | (Vincent, 2017) |

**Table S2.** Sample preparation, instrumental analysis, calibration, and quality-control procedures used for mandibular elemental quantification.

| Analytical stage | Procedure or specification |
|---|---|
| Analytical sample | Left and right mandibles from each individual combined as a single analytical sample |
| Initial tissue preparation | Heads processed in deionised water; surrounding soft tissues mechanically removed using ceramic and plastic tools |
| Cleaning | Mandibles rinsed with deionised water and placed in an ultrasonic bath |
| Drying | Oven-dried at 60°C for 48 h |
| Tooth removal | All teeth removed using forceps before digestion |
| Visual inspection | Each mandible inspected to confirm the absence of residual soft tissues |
| Sample weighing | Combined mandibular sample weighed to the nearest 0.1 mg using an AS 220.R2 PLUS analytical balance (RADWAG) |
| Storage | Samples stored individually in polypropylene vials under dry and dark laboratory conditions |
| Digestion vessel | Closed Teflon vessel |
| First digestion step | 2 mL concentrated $HNO_3$; heated at 75°C for 2 h |
| Second digestion step | After cooling, 0.5 mL $H_2O_2$ added; samples left overnight at room temperature |
| Evaporation and reconstitution | Samples evaporated at 100°C and residues redissolved in 5 mL of 2% $HNO_3$ |
| ICP-OES instrument | iCAP 7000 series, Thermo Scientific, Germany |
| Elements quantified by ICP-OES | Ca, Mg, Na, K, Zn, Fe, Pb, Al, Sr, and Ba |
| ICP-MS instrument | iCAP Q, Thermo Scientific, Germany |
| Elements quantified by ICP-MS | As, B, Co, Cr, Cu, Li, Mn, Mo, Ni, and V |
| ICP-OES calibration standard | AN9090MN, CRM Mix 26 elements, Analytika |
| ICP-MS calibration standard | ICP multi-element standard solution VI, CRM Mix 30 elements, Merck |
| Quality control | Procedural blanks, duplicate samples, and certified reference materials |
| Certified reference materials | NIST SRM 1643f, Trace Elements in Water; NIST SRM 1400, Bone Ash |
| Reporting units | mg $kg^{-1}$ (ppm) |
| Analytical laboratory | Department of Geosciences, Czech University of Life Sciences Prague, Czechia |

**Table S3.** Sample sizes by species, season, sex, and habitat type.

| Factor | Category | *A. flavicollis* individuals | *C. glareolus* individuals | Total (n) |
|---|---|---|---|---|
| **Season** | Spring | 29 | 21 | 50 |
| | Autumn | 16 | 18 | 34 |
| **Sex** | Female | 22 | 20 | 42 |
| | Male | 23 | 19 | 42 |
| **Habitat type** | Spruce forest | 20 | 11 | 31 |
| | Beech forest | 25 | 28 | 53 |
| **Total** | | **45** | **39** | **84** |

**Table S4.** Descriptive statistics of mandibular elemental concentrations in *Apodemus flavicollis* and *Clethrionomys glareolus*.

| Category | Element | *Apodemus flavicollis* | | | *Clethrionomys glareolus* | | |
|---|---|---|---|---|---|---|---|
| | | **Mean ± SD** | **Min** | **Max** | **Mean ± SD** | **Min** | **Max** |
| **Essential** | Ca | 265157.70 ± 25131.21 | 224450.3 | 335269.6 | 273183.70 ± 22921.74 | 230560 | 339584.8 |
| | Co | 1.01 ± 0.11 | 0.81 | 1.21 | 1.02 ± 0.16 | 0.45 | 1.32 |
| | Cu | 0.67 ± 0.82 | 0.09 | 4.82 | 0.83 ± 0.79 | 0.05 | 3.98 |
| | Fe | 216.32 ± 59.48 | 117.72 | 350.26 | 275.50 ± 80.38 | 176.25 | 561.66 |
| | K | 42.15 ± 22.40 | 9.75 | 125.57 | 35.35 ± 16.31 | 15.8 | 91.62 |
| | Mg | 2436.38 ± 613.02 | 1100.95 | 3779.43 | 2257.59 ± 612.41 | 1049.42 | 3666.33 |
| | Mn | 2.62 ± 1.42 | 0.8 | 7.16 | 7.22 ± 4.64 | 2.18 | 22.87 |
| | Mo | 0.07 ± 0.12 | 0.02 | 0.69 | 0.10 ± 0.15 | 0.02 | 0.91 |
| | Na | 2388.59 ± 609.57 | 1102.95 | 3489 | 2142.18 ± 453.09 | 1121.83 | 3651.45 |
| | Zn | 311.95 ± 62.13 | 182.24 | 535.48 | 338.68 ± 43.57 | 257.2 | 458.12 |
| **Non-essential** | Al | 14.71 ± 4.55 | 8.79 | 33.82 | 17.88 ± 6.32 | 10.35 | 35.8 |
| | As | 1.36 ± 0.31 | 0.42 | 2.12 | 1.21 ± 0.46 | 0.4 | 2.23 |
| | B | 2.34 ± 0.88 | 1 | 4.37 | 3.31 ± 3.29 | 1.22 | 15.97 |
| | Ba | 356.14 ± 162.72 | 80.24 | 831.36 | 319.78 ± 108.44 | 149.86 | 584.42 |
| | Cr | 0.58 ± 0.32 | 0.21 | 1.33 | 0.55 ± 0.27 | 0.11 | 1.27 |
| | Li | 0.12 ± 0.09 | 0.05 | 0.4 | 0.15 ± 0.08 | 0.04 | 0.36 |
| | Ni | 6.78 ± 0.89 | 4.9 | 8.53 | 6.99 ± 1.38 | 2.17 | 9.66 |
| | Pb | 17.47 ± 4.85 | 6.48 | 29.57 | 21.34 ± 7.99 | 5.43 | 35.49 |
| | Sr | 335.25 ± 118.45 | 96.44 | 604.75 | 237.12 ± 95.82 | 59.26 | 566.21 |
| | V | 0.62 ± 0.28 | 0.29 | 1.65 | 1.35 ± 1.90 | 0.26 | 8.29 |

**Note.** Values are presented as mean ± sd, minimum, and maximum. Sample sizes were *A. flavicollis* (n = 45) and *C. glareolus* (n = 39). Elemental concentrations are expressed in mg $kg^{-1}$ (ppm).

**Table S5.** Relative contributions of biological, environmental, interactions, and residual components of essential and non-essential mandibular elementomes in the pooled dataset and separately for each species.

| Dataset | Element group | Component | $R^2$ | Variation explained (%) |
|---|---|---|---|---|
| Both species | Essential | Biological | 0.1968 | 19.7 |
| | | Environmental | 0.1185 | 11.8 |
| | | Interactions | 0.0707 | 7.1 |
| | | Residuals | 0.6141 | 61.4 |
| | Non-essential | Biological | 0.1331 | 13.3 |
| | | Environmental | 0.1828 | 18.3 |
| | | Interactions | 0.0914 | 9.1 |
| | | Residuals | 0.5927 | 59.3 |
| *A. flavicollis* | Essential | Biological | 0.1215 | 12.2 |
| | | Environmental | 0.1079 | 10.8 |
| | | Interactions | 0.0842 | 8.4 |
| | | Residuals | 0.6863 | 68.6 |
| | Non-essential | Biological | 0.0799 | 8.0 |
| | | Environmental | 0.2163 | 21.6 |
| | | Interactions | 0.1220 | 12.2 |
| | | Residuals | 0.5818 | 58.2 |
| *C. glareolus* | Essential | Biological | 0.0644 | 6.4 |
| | | Environmental | 0.1603 | 16.0 |
| | | Interactions | 0.0752 | 7.5 |
| | | Residuals | 0.7002 | 70.0 |
| | Non-essential | Biological | 0.0699 | 7.0 |
| | | Environmental | 0.1809 | 18.1 |
| | | Interactions | 0.0811 | 8.1 |
| | | Residuals | 0.6682 | 66.8 |

**Note.** Values were obtained from the $R^2$ contributions of terms in the PERMANOVA models. In the pooled analyses, the biological component included species, body mass × species, and sex × species, whereas season × species and habitat type × species were part of the environmental component. In species-specific analyses, biological predictors included body mass and sex, whereas season and habitat type were part of the environmental component. The interaction component comprised the remaining interaction terms.

**Table S6.** PERMANOVA results for *clr*-transformed essential and non-essential mandibular elementomes in the complete dataset and separately for each species.

| | Element group | Predictor | df | F | $R^2$ | *p*-value |
|---|---|---|---|---|---|---|
| **Both species** | Essential | Species | 1 | 12.26 | 0.114 | 0.001 |
| | | Body mass × Species | 2 | 4.23 | 0.079 | 0.002 |
| | | Season × Species | 2 | 4.37 | 0.081 | 0.002 |
| | | Sex × Species | 2 | 0.22 | 0.004 | 0.992 |
| | | Habitat type × Species | 2 | 2.00 | 0.037 | 0.048 |
| | | Body mass × Season × Species | 2 | 2.53 | 0.047 | 0.020 |
| | | Body mass × Sex × Species | 2 | 0.67 | 0.013 | 0.674 |
| | | Season × Habitat type × Species | 2 | 0.52 | 0.010 | 0.845 |
| | | Body mass × Habitat type × Species | 2 | 0.08 | 0.001 | 1.000 |
| | Non-essential | Species | 1 | 7.11 | 0.064 | 0.001 |
| | | Body mass × Species | 2 | 3.22 | 0.058 | 0.002 |
| | | Season × Species | 2 | 5.70 | 0.102 | 0.001 |
| | | Sex × Species | 2 | 0.63 | 0.011 | 0.816 |
| | | Habitat type × Species | 2 | 4.47 | 0.080 | 0.001 |
| | | Body mass × Season × Species | 2 | 2.57 | 0.046 | 0.008 |
| | | Body mass × Sex × Species | 2 | 0.41 | 0.007 | 0.960 |
| | | Season × Habitat type × Species | 2 | 1.60 | 0.029 | 0.096 |
| | | Body mass × Habitat type × Species | 2 | 0.50 | 0.009 | 0.928 |
| ***Apodemus flavicollis*** | Essential | Body mass | 1 | 6.08 | 0.116 | 0.002 |
| | | Season | 1 | 4.86 | 0.093 | 0.004 |
| | | Sex | 1 | 0.3 | 0.006 | 0.9 |
| | | Habitat type | 1 | 0.8 | 0.015 | 0.506 |
| | | Body mass × Season | 1 | 2.2 | 0.042 | 0.081 |
| | | Body mass × Sex | 1 | 1.15 | 0.022 | 0.326 |
| | | Season × Habitat type | 1 | 0.99 | 0.019 | 0.399 |
| | | Body mass × Habitat type | 1 | 0.08 | 0.001 | 0.999 |
| | Non-essential | Body mass | 1 | 4.52 | 0.073 | 0.003 |
| | | Season | 1 | 8.82 | 0.143 | 0.001 |
| | | Sex | 1 | 0.42 | 0.007 | 0.834 |
| | | Habitat type | 1 | 4.56 | 0.074 | 0.002 |
| | | Body mass × Season | 1 | 4.13 | 0.067 | 0.009 |
| | | Body mass × Sex | 1 | 0.33 | 0.005 | 0.905 |
| | | Season × Habitat type | 1 | 2.84 | 0.046 | 0.035 |
| | | Body mass × Habitat type | 1 | 0.24 | 0.004 | 0.949 |
| ***Clethrionomys glareolus*** | Essential | Body mass | 1 | 2.61 | 0.061 | 0.065 |
| | | Season | 1 | 3.89 | 0.091 | 0.013 |
| | | Sex | 1 | 0.15 | 0.004 | 0.972 |
| | | Habitat type | 1 | 2.98 | 0.07 | 0.03 |
| | | Body mass × Season | 1 | 2.77 | 0.065 | 0.046 |
| | | Body mass × Sex | 1 | 0.27 | 0.006 | 0.897 |
| | | Season × Habitat type | 1 | 0.12 | 0.003 | 0.987 |
| | | Body mass × Habitat type | 1 | 0.07 | 0.002 | 0.993 |
| | Non-essential | Body mass | 1 | 2.43 | 0.054 | 0.04 |
| | | Season | 1 | 3.9 | 0.087 | 0.003 |
| | | Sex | 1 | 0.7 | 0.016 | 0.652 |
| | | Habitat type | 1 | 4.22 | 0.094 | 0.004 |
| | | Body mass × Season | 1 | 1.68 | 0.037 | 0.136 |
| | | Body mass × Sex | 1 | 0.44 | 0.01 | 0.871 |
| | | Season × Habitat type | 1 | 0.91 | 0.02 | 0.427 |
| | | Body mass × Habitat type | 1 | 0.61 | 0.014 | 0.725 |

**Note.** Pooled-species models included species, species-specific effects of body mass, season, sex, and habitat type, and the specified three-way interactions. Species-specific models excluded species. Sequential effects were tested using 999 permutations. $R^2$ represents the sequential proportion of multivariate variation associated with each model term.

**Table S7.** Comparison of interspecific differentiation in multielemental and Ba/Ca–Sr/Ca ratio spaces.

| **Metric** | **Multielemental niche** | **Ba/Ca–Sr/Ca ratio space** |
|---|---|---|
| Dimensions | PC1–PC5 | log10(Ba/Ca)–log10(Sr/Ca) |
| Cumulative variance (%) | 74.8 | — |
| PERMANOVA ($F_{1,82}$) | 10.485 | 9.431 |
| PERMANOVA ($R^2$) | 0.113 | 0.103 |
| PERMANOVA (p) | 0.001 | 0.002 |
| PERMDISP ($F_{1,82}$) | 0.615 | 0.748 |
| PERMDISP (p) | 0.444 | 0.393 |
| Centroid distance | 1.168 | 0.169 |
| Convex-hull overlap relative to union (%) | 34.64 | 39.46 |
| Mean distance to centroid: *A. flavicollis* **individuals** | 1.467 | 0.208 |
| Mean distance to centroid: *C. glareolus* **individuals** | 1.574 | 0.178 |

**Note.** Multi-elemental centroid separation and within-species dispersion were calculated in the retained PC1–PC5 space, whereas convex-hull overlap was calculated in the PC1–PC2 projection. All ratio-based metrics were calculated in the two-dimensional $\log_{10}$(Ba/Ca)–$\log_{10}$(Sr/Ca) space. Because the representations differ in dimensionality and scale, absolute centroid and dispersion distances should not be compared directly between them. The two distance matrices were weakly but significantly correlated (Pearson Mantel test $r = 0.23$, p = 0.003; 999 permutations).

**Table S8.** Significant body-mass allometric relationships from element-specific linear models of mandibular elemental concentrations in *Apodemus flavicollis* and *Clethrionomys glareolus*.

| Type | Element | Scaling *b* | SE | *p value* |
|---|---|---|---|---|
| | | ***A. flavicollis*** | | |
| Essential | Ca | –0.155 | 0.041 | 0.0005 |
| Essential | Cu | –1.022 | 0.437 | 0.0243 |
| Essential | Mg | 0.716 | 0.118 | < 0.0001 |
| Essential | Mo | –1.006 | 0.419 | 0.021 |
| Essential | Na | 0.712 | 0.103 | < 0.0001 |
| Non-essential | Al | –0.668 | 0.118 | < 0.0001 |
| Non-essential | Cr | –0.705 | 0.167 | 0.0001 |
| Non-essential | V | –0.739 | 0.147 | < 0.0001 |
| | | ***C. glareolus*** | | |
| Essential | Mg | 0.731 | 0.184 | 0.0003 |
| Essential | Mn | –1.312 | 0.445 | 0.0057 |
| Essential | Na | 0.46 | 0.147 | 0.0036 |

**Note.** Elemental concentrations and body mass were $\log_{10}$-transformed before analysis. Models included body mass, season, and habitat type as predictors. Estimates (scalings *b*), standard errors (SE), and p-values are reported. Only statistically significant body-mass effects are shown.

**Table S9**. Robust allometric relationships between body mass and mandibular concentrations of Pb, Sr, Ba, Mg, and Ca, stratified by sex and season, in *Apodemus flavicollis* and *Clethrionomys glareolus*.

| Element | Species | Group | n | Scaling $b_i$ | SE | t | Approx. p-value | 95% CI |
|---|---|---|---|---|---|---|---|---|
| | | | | **Panel A.** Sex-specific slopes | | | | |
| Pb | *A. flavicollis* | Female | 22 | 0.135 | 0.251 | 0.539 | 0.590 | (−0.212, 0.640) |
| Pb | *A. flavicollis* | Male | 23 | –0.367 | 0.180 | –2.038 | 0.042 | (−0.839, 0.039) |
| Pb | *C. glareolus* | Female | 20 | 0.603 | 0.392 | 1.539 | 0.124 | (−0.499, 1.644) |
| Pb | *C. glareolus* | Male | 19 | 0.247 | 0.487 | 0.507 | 0.612 | (−0.988, 1.066) |
| Sr | *A. flavicollis* | Female | 22 | 0.501 | 0.375 | 1.337 | 0.181 | (−0.354, 1.589) |
| Sr | *A. flavicollis* | Male | 23 | –0.190 | 0.228 | -0.833 | 0.405 | (−0.673, 0.133) |
| Sr | *C. glareolus* | Female | 20 | 0.040 | 0.491 | 0.082 | 0.935 | (−0.695, 1.145) |
| Sr | *C. glareolus* | Male | 19 | –0.926 | 0.316 | –2.934 | 0.003 | (−1.387, −0.178) |
| Ba | *A. flavicollis* | Female | 22 | 0.893 | 0.493 | 1.809 | 0.070 | (−0.610, 2.159) |
| Ba | *A. flavicollis* | Male | 23 | –0.275 | 0.295 | –0.933 | 0.351 | (−0.910, 0.088) |
| Ba | *C. glareolus* | Female | 20 | –0.095 | 0.366 | –0.261 | 0.794 | (−0.860, 0.805) |
| Ba | *C. glareolus* | Male | 19 | –0.703 | 0.337 | –2.090 | 0.037 | (−1.262, 0.438) |
| Mg | *A. flavicollis* | Female | 22 | 0.644 | 0.140 | 4.605 | <0.001 | (0.265, 0.843) |
| Mg | *A. flavicollis* | Male | 23 | 0.829 | 0.140 | 5.940 | <0.001 | (0.449, 1.109) |
| Mg | *C. glareolus* | Female | 20 | 0.655 | 0.226 | 2.896 | 0.004 | (0.061, 1.477) |
| Mg | *C. glareolus* | Male | 19 | 1.268 | 0.236 | 5.365 | <0.001 | (0.101, 1.979) |
| Ca | *A. flavicollis* | Female | 22 | –0.149 | 0.077 | –1.933 | 0.053 | (−0.322, −0.001) |
| Ca | *A. flavicollis* | Male | 23 | –0.189 | 0.063 | –2.997 | 0.003 | (−0.308, −0.035) |
| Ca | *C. glareolus* | Female | 20 | –0.082 | 0.109 | –0.752 | 0.452 | (−0.380, 0.097) |
| Ca | *C. glareolus* | Male | 19 | 0.019 | 0.101 | 0.191 | 0.849 | (−0.319, 0.174) |
| | | | | **Panel B**. Season-specific slopes | | | | |
| Pb | *A. flavicollis* | Autumn | 16 | –0.010 | 0.347 | –0.028 | 0.978 | (−1.013, 0.550) |
| Pb | *A. flavicollis* | Spring | 29 | –0.177 | 0.189 | –0.935 | 0.350 | (−0.416, 0.150) |
| Pb | *C. glareolus* | Autumn | 18 | –0.728 | 0.442 | –1.648 | 0.100 | (−1.431, 0.241) |
| Pb | *C. glareolus* | Spring | 21 | 1.116 | 0.377 | 2.961 | 0.003 | (0.571, 2.171) |
| Sr | *A. flavicollis* | Autumn | 16 | 1.105 | 0.524 | 2.106 | 0.035 | (0.494, 3.075) |
| Sr | *A. flavicollis* | Spring | 29 | –0.150 | 0.167 | –0.898 | 0.369 | (−0.413, 0.118) |
| Sr | *C. glareolus* | Autumn | 18 | –0.752 | 0.359 | –2.093 | 0.036 | (−1.317, −0.225) |
| Sr | *C. glareolus* | Spring | 21 | –0.401 | 0.369 | –1.085 | 0.278 | (−1.179, 1.106) |
| Ba | *A. flavicollis* | Autumn | 16 | 1.919 | 0.613 | 3.130 | 0.002 | (1.136, 3.211) |
| Ba | *A. flavicollis* | Spring | 29 | –0.214 | 0.267 | –0.799 | 0.425 | (−0.714, 0.159) |
| Ba | *C. glareolus* | Autumn | 18 | 0.281 | 0.428 | 0.655 | 0.512 | (−0.479, 0.948) |
| Ba | *C. glareolus* | Spring | 21 | –0.919 | 0.295 | –3.114 | 0.002 | (−1.464, −0.169) |
| Mg | *A. flavicollis* | Autumn | 16 | 0.769 | 0.164 | 4.674 | <0.001 | (0.337, 1.589) |
| Mg | *A. flavicollis* | Spring | 29 | 0.703 | 0.128 | 5.488 | <0.001 | (0.319, 0.957) |
| Mg | *C. glareolus* | Autumn | 18 | 1.769 | 0.202 | 8.772 | <0.001 | (1.150, 2.053) |
| Mg | *C. glareolus* | Spring | 21 | 0.170 | 0.218 | 0.777 | 0.437 | (−0.247, 0.738) |
| Ca | *A. flavicollis* | Autumn | 16 | –0.155 | 0.066 | –2.334 | 0.020 | (−0.253, 0.006) |
| Ca | *A. flavicollis* | Spring | 29 | –0.164 | 0.049 | –3.367 | <0.001 | (−0.269, −0.034) |
| Ca | *C. glareolus* | Autumn | 18 | –0.135 | 0.110 | –1.233 | 0.217 | (−0.349, 0.035) |
| Ca | *C. glareolus* | Spring | 21 | –0.029 | 0.067 | –0.439 | 0.661 | (−0.269, 0.092) |

**Note.** Slopes were estimated using robust linear models fitted to $\log_{10}$-transformed body mass and elemental concentrations. Approximate *p*-values are based on a two-sided normal approximation. The 95% confidence intervals for the slopes were obtained using percentile bootstrap with 999 resamples.

**Table S10.** Robust allometric relationships between body mass and mandibular Pb/Ca, Sr/Ca, Ba/Ca, and Mg/Ca ratios, stratified by sex and season, in *Apodemus flavicollis* and *Clethrionomys glareolus.*

| Element | Species | Group | n | Scaling $b_{ratio}$ | SE | t | Approx. p-value | 95% CI |
|---|---|---|---|---|---|---|---|---|
| **Panel A.** Sex-specific slopes | | | | | | | | |
| Pb/Ca | *A. flavicollis* | Female | 22 | 0.321 | 0.288 | 1.114 | 0.265 | (−0.101, 0.792) |
| Pb/Ca | *A. flavicollis* | Male | 23 | –0.194 | 0.189 | –1.024 | 0.306 | (−0.623, 0.217) |
| Pb/Ca | *C. glareolus* | Female | 20 | 0.848 | 0.498 | 1.703 | 0.089 | (−0.391, 2.104) |
| Pb/Ca | *C. glareolus* | Male | 19 | 0.227 | 0.462 | 0.491 | 0.623 | (−0.673, 1.108) |
| Sr/Ca | *A. flavicollis* | Female | 22 | 0.615 | 0.350 | 1.755 | 0.079 | (−0.059, 1.537) |
| Sr/Ca | *A. flavicollis* | Male | 23 | –0.040 | 0.201 | -0.197 | 0.844 | (−0.389, 0.254) |
| Sr/Ca | *C. glareolus* | Female | 20 | 0.131 | 0.397 | 0.330 | 0.742 | (−0.420, 0.999) |
| Sr/Ca | *C. glareolus* | Male | 19 | –0.939 | 0.298 | –3.150 | 0.002 | (−1.329, −0.069) |
| Ba/Ca | *A. flavicollis* | Female | 22 | 0.882 | 0.472 | 1.869 | 0.062 | (−0.389, 2.139) |
| Ba/Ca | *A. flavicollis* | Male | 23 | –0.092 | 0.272 | –0.339 | 0.735 | (−0.760, 0.284) |
| Ba/Ca | *C. glareolus* | Female | 20 | 0.054 | 0.345 | 0.157 | 0.875 | (−0.756, 0.744) |
| Ba/Ca | *C. glareolus* | Male | 19 | –0.780 | 0.343 | –2.274 | 0.023 | (−1.329, 0.486) |
| Mg/Ca | *A. flavicollis* | Female | 22 | 0.846 | 0.127 | 6.648 | <0.001 | (0.392, 1.099) |
| Mg/Ca | *A. flavicollis* | Male | 23 | 1.080 | 0.123 | 8.779 | <0.001 | (0.683, 1.357) |
| Mg/Ca | *C. glareolus* | Female | 20 | 0.794 | 0.258 | 3.073 | 0.002 | (0.213, 1.582) |
| Mg/Ca | *C. glareolus* | Male | 19 | 1.356 | 0.255 | 5.312 | <0.001 | (−0.071, 2.073) |
| **Panel B.** Season-specific slopes | | | | | | | | |
| Pb/Ca | *A. flavicollis* | Autumn | 16 | 0.110 | 0.383 | 0.286 | 0.775 | (−0.983, 0.675) |
| Pb/Ca | *A. flavicollis* | Spring | 29 | 0.004 | 0.187 | 0.020 | 0.984 | (−0.273, 0.354) |
| Pb/Ca | *C. glareolus* | Autumn | 18 | –0.530 | 0.364 | –1.456 | 0.145 | (−1.184, 0.381) |
| Pb/Ca | *C. glareolus* | Spring | 21 | 1.180 | 0.428 | 2.761 | 0.006 | (0.624, 2.435) |
| Sr/Ca | *A. flavicollis* | Autumn | 16 | 1.244 | 0.503 | 2.474 | 0.013 | (0.685, 3.048) |
| Sr/Ca | *A. flavicollis* | Spring | 29 | –0.027 | 0.143 | –0.189 | 0.850 | (−0.212, 0.269) |
| Sr/Ca | *C. glareolus* | Autumn | 18 | –0.628 | 0.391 | –1.606 | 0.108 | (−1.206, −0.078) |
| Sr/Ca | *C. glareolus* | Spring | 21 | –0.169 | 0.310 | –0.544 | 0.587 | (−1.254, 1.328) |
| Ba/Ca | *A. flavicollis* | Autumn | 16 | 2.061 | 0.483 | 4.266 | <0.001 | (1.261, 3.316) |
| Ba/Ca | *A. flavicollis* | Spring | 29 | –0.088 | 0.262 | –0.335 | 0.737 | (−0.504, 0.304) |
| Ba/Ca | *C. glareolus* | Autumn | 18 | 0.414 | 0.440 | 0.939 | 0.348 | (−0.462, 1.135) |
| Ba/Ca | *C. glareolus* | Spring | 21 | –0.751 | 0.299 | –2.511 | 0.012 | (−1.529, −0.023) |
| Mg/Ca | *A. flavicollis* | Autumn | 16 | 0.881 | 0.167 | 5.264 | <0.001 | (0.377, 1.849) |
| Mg/Ca | *A. flavicollis* | Spring | 29 | 0.966 | 0.147 | 6.593 | <0.001 | (0.545, 1.174) |
| Mg/Ca | *C. glareolus* | Autumn | 18 | 1.921 | 0.153 | 12.525 | <0.001 | (1.421, 2.135) |
| Mg/Ca | *C. glareolus* | Spring | 21 | 0.219 | 0.235 | 0.934 | 0.350 | (−0.228, 0.831) |

**Note**: Slopes were estimated using robust linear models fitted to $\log_{10}$-transformed body mass and elemental ratios. Approximate *p*-values are based on a two-sided normal approximation. The 95% confidence intervals for the slopes were obtained using percentile bootstrap with 999 resamples.

**Table S11.** Significant seasonal and habitat effects from element-specific linear models of mandibular elemental concentrations in *Apodemus flavicollis* and *Clethrionomys glareolus*.

| Type | Element | Predictor | Term | β | SE | *p value* |
|---|---|---|---|---|---|---|
| | | | ***A. flavicollis*** | | | |
| Essential | Fe | Habitat type | Beech forest | 0.105 | 0.035 | 0.0044 |
| Essential | Ca | Season | Spring | 0.027 | 0.01 | 0.0115 |
| Essential | Co | Season | Spring | 0.044 | 0.014 | 0.0038 |
| Essential | Mn | Season | Spring | 0.245 | 0.062 | 0.0003 |
| Non-essential | Al | Season | Spring | 0.06 | 0.029 | 0.0449 |
| Non-essential | As | Season | Spring | 0.098 | 0.034 | 0.0057 |
| Non-essential | B | Season | Spring | 0.291 | 0.03 | < 0.0001 |
| Non-essential | Ba | Season | Spring | 0.193 | 0.062 | 0.0033 |
| Non-essential | Cr | Season | Spring | 0.359 | 0.041 | < 0.0001 |
| Non-essential | Li | Season | Spring | 0.209 | 0.076 | 0.0086 |
| Non-essential | Ni | Season | Spring | 0.042 | 0.018 | 0.0258 |
| Non-essential | Sr | Season | Spring | 0.139 | 0.049 | 0.0074 |
| Non-essential | V | Season | Spring | 0.207 | 0.036 | < 0.0001 |
| | | | ***C. glareolus*** | | | |
| Essential | Mg | Habitat type | Beech forest | 0.11 | 0.037 | 0.0055 |
| Essential | Na | Habitat type | Beech forest | 0.081 | 0.03 | 0.0098 |
| Non-essential | Al | Habitat type | Beech forest | –0.142 | 0.04 | 0.0012 |
| Non-essential | Ba | Habitat type | Beech forest | 0.135 | 0.048 | 0.0079 |
| Non-essential | Sr | Habitat type | Beech forest | 0.151 | 0.059 | 0.0144 |
| Non-essential | V | Habitat type | Beech forest | –0.37 | 0.12 | 0.004 |
| Essential | Ca | Season | Spring | 0.035 | 0.012 | 0.005 |
| Non-essential | As | Season | Spring | 0.189 | 0.068 | 0.0085 |
| Non-essential | Cr | Season | Spring | 0.324 | 0.066 | 0 |

**Note.** Elemental concentrations and body mass were $\log_{10}$-transformed before analysis. Models included body mass, season, and habitat type as predictors. Estimates (β), standard errors (SE), and p-values are reported. Autumn and spruce forest were used as reference levels for season and habitat type, respectively. Positive and negative β values indicate higher and lower elemental concentrations in the "term" category, respectively, relative to the corresponding reference category. Only statistically significant seasonal and habitat effects are shown.

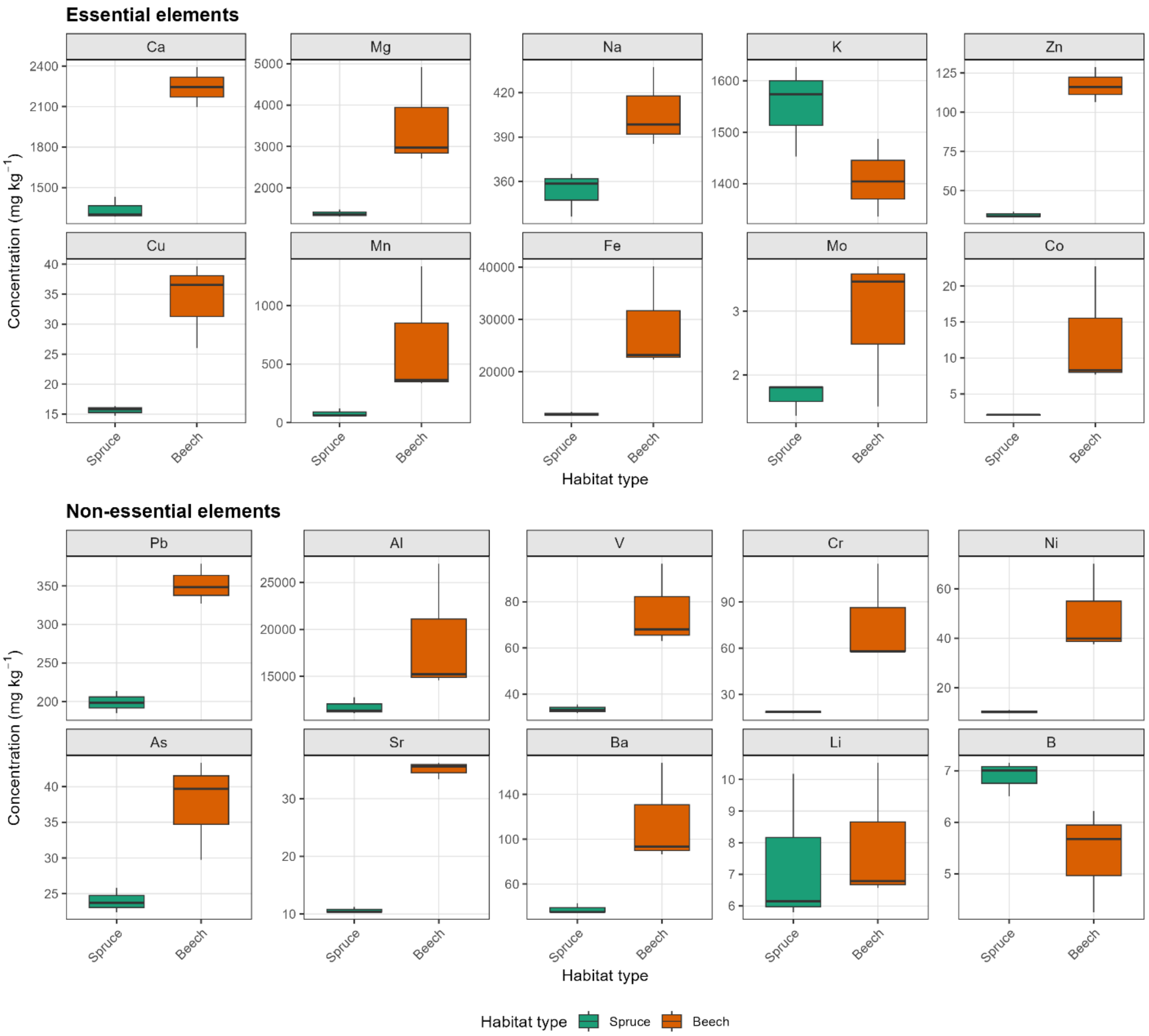


**Figure S1.** Soil elemental concentrations in spruce- and beech-dominated forest habitats in the Ore Mountains, Czechia. Boxplots show concentrations of essential and non-essential elements in soil samples collected from the two habitat types (n=3 per habitat type). Colours indicate habitat type (spruce forest and beech forest). Elemental concentrations are expressed in mg kg$^{-1}$.

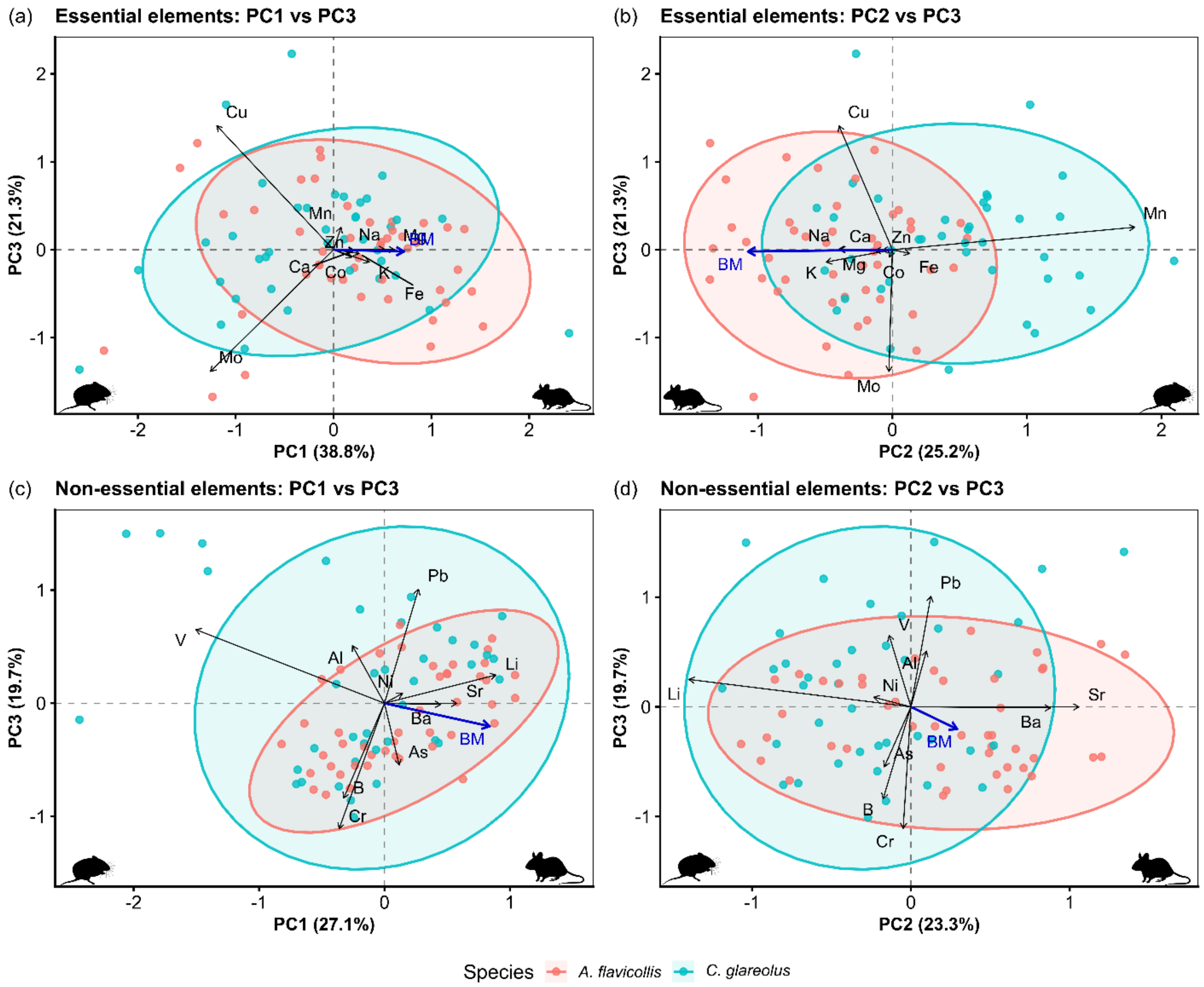


**Figure S2.** Principal component analysis (PCA) Biplots including the third axis for essential and non-essential mandibular elementomes. PCAs were conducted on *clr*-transformed elemental compositions. Panels show PC1 vs. PC3 and PC2 vs. PC3 biplots for essential elements (a–b) and non-essential elements (c–d). The first three principal components explained 85.2% and 70.1% of the cumulative variance in essential and non-essential elementomes, respectively. Points represent individuals, colours show species, and ellipses represent 95% ellipses for each species. Black arrows indicate element loadings, and the blue vector shows the significant association of body mass with the ordination, fitted as a supplementary variable. Animal silhouettes were obtained from 'PhyloPic.org' (public domain).

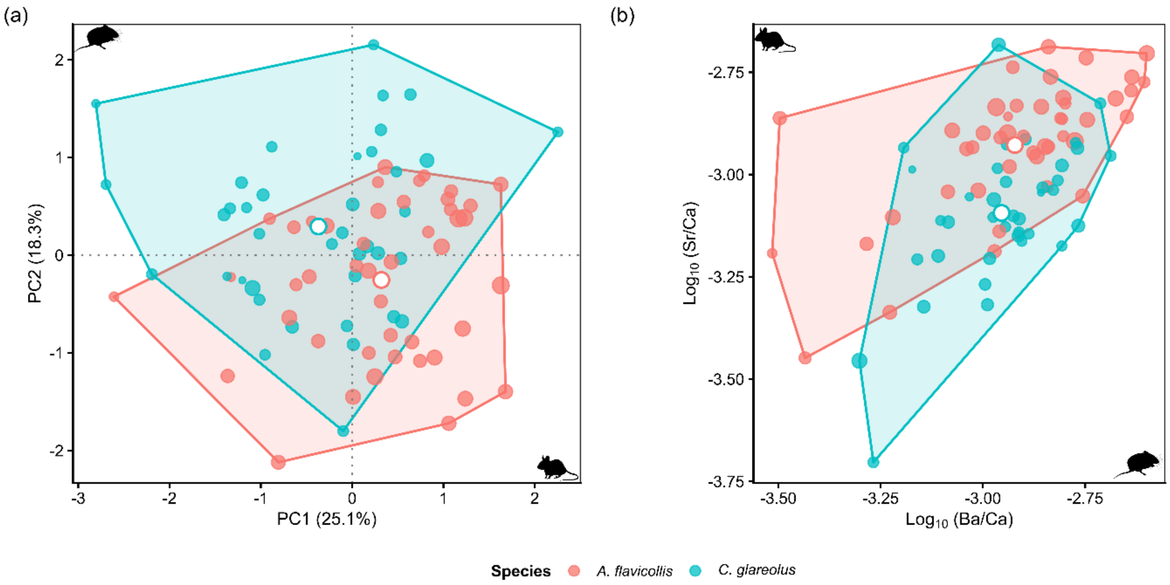


**Figure S3**. Comparison of interspecific elemental differentiation in multi-elemental and ratio-based spaces. Panel (a) shows the PC1–PC2 projection of the *clr*-transformed complete elementome, whereas panel (b) shows the space defined by $\log_{10}$-transformed Ba/Ca and Sr/Ca ratios. Points represent individuals, point size is proportional to body mass, shaded polygons represent species-specific convex hulls, open circles indicate species centroids, and colours distinguish species. Animal silhouettes were obtained from 'PhyloPic.org' (public domain).

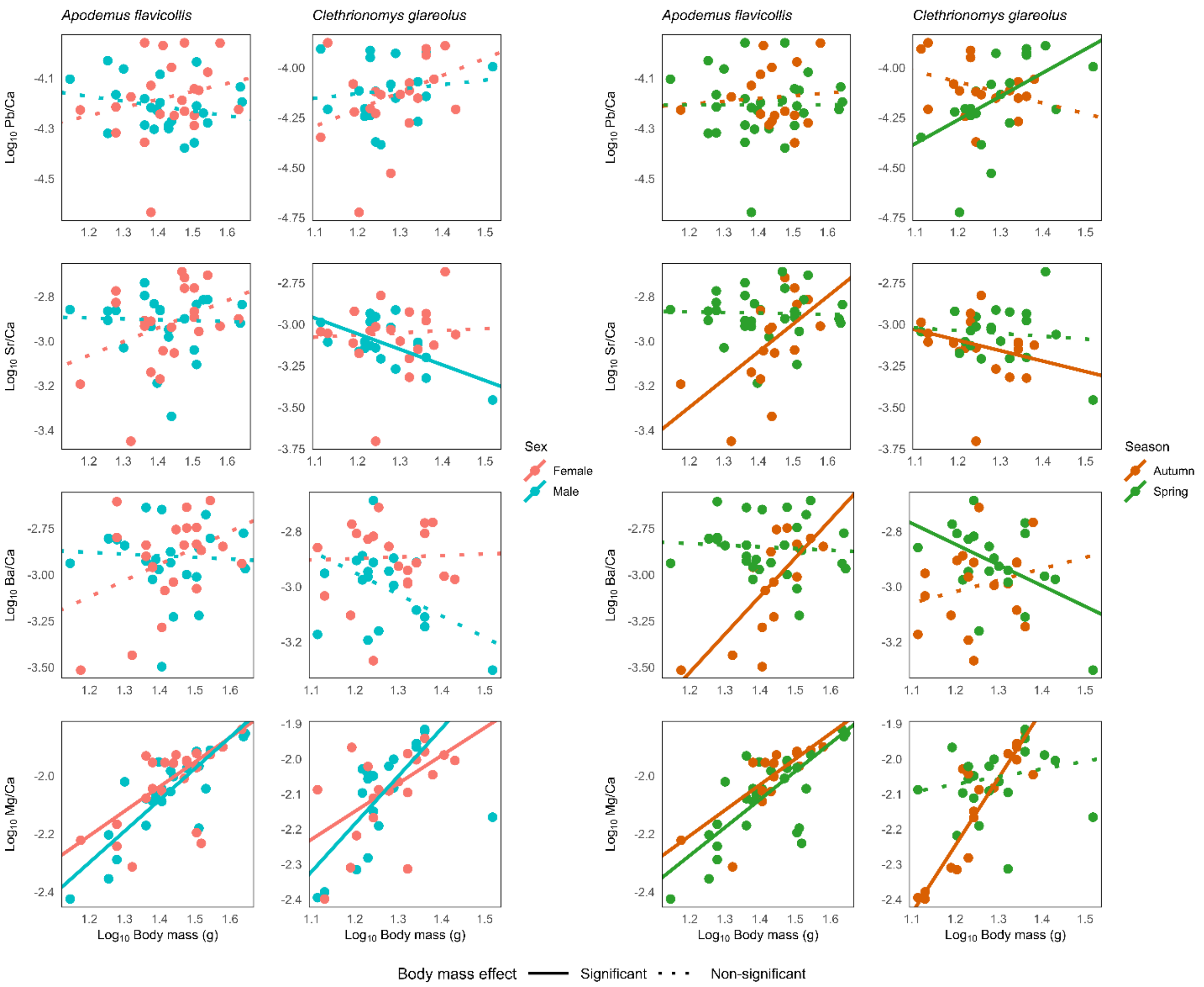


**Figure S4.** Log-log relationships between body mass and mandibular elemental ratios in *Apodemus flavicollis* and *Clethrionomys glareolus*, stratified by (a) sex and (b) season. Panels show Pb/Ca, Sr/Ca, Ba/Ca, and Mg/Ca relationships. Points represent individual observations, and lines show group-specific robust linear regressions. Solid and dotted lines indicate significant and non-significant body-mass effects, respectively. Robust slope estimates (allometric scalings *b*), bootstrap confidence intervals, and associated statistics are provided in Table S10.

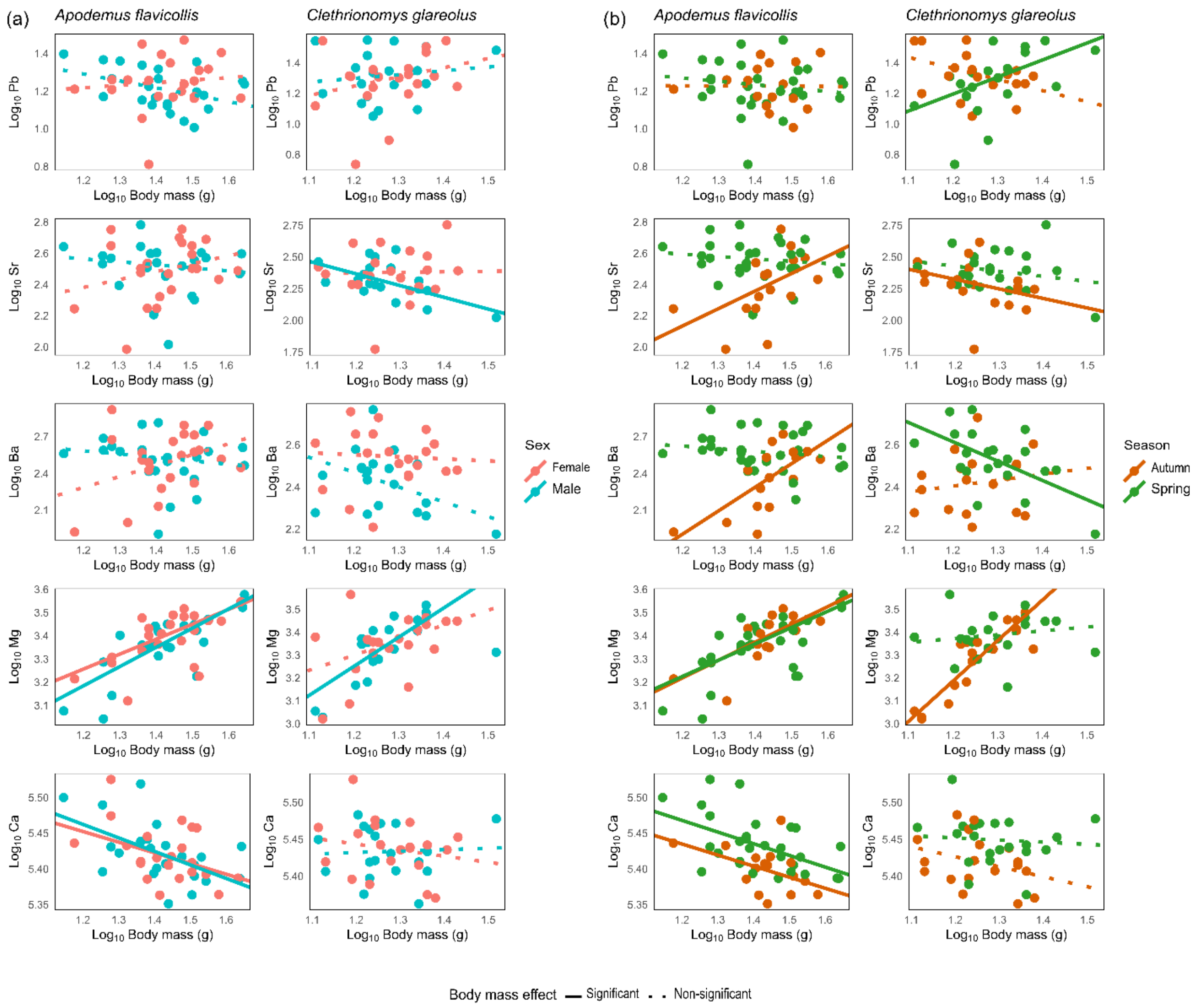


**Figure S5**. Allometric relationships between body mass and mandibular elemental concentrations in *Apodemus flavicollis* and *Clethrionomys glareolus*. Panels show relationships for Pb, Sr, Ba, Mg, and Ca, stratified by sex (left-hand panels) and season (right-hand panels). Points represent individual observations, and lines show group-specific robust linear regressions. Solid and dotted lines indicate significant and non-significant body-mass effects, respectively. Both axes are $log_{10}$-transformed. Robust scaling estimates, bootstrap confidence intervals, and associated statistics are provided in Table S9.

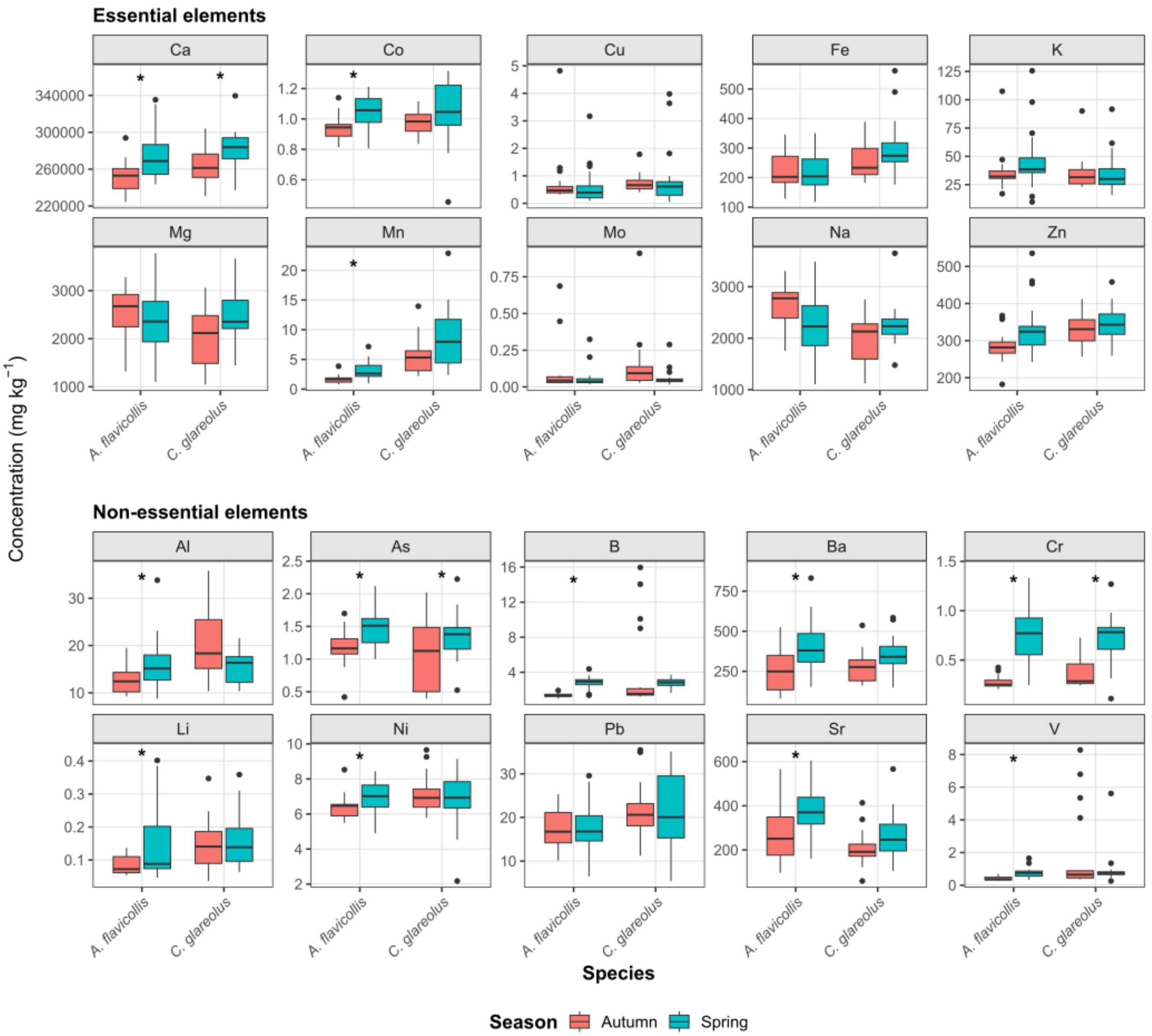


**Figure S6.** Seasonal variation in mandibular elemental concentrations of *Apodemus flavicollis* and *Clethrionomys glareolus*. Boxplots show essential and non-essential elemental concentrations by season within each species. Asterisks indicate significant seasonal effects from element-specific linear models (Table S11).

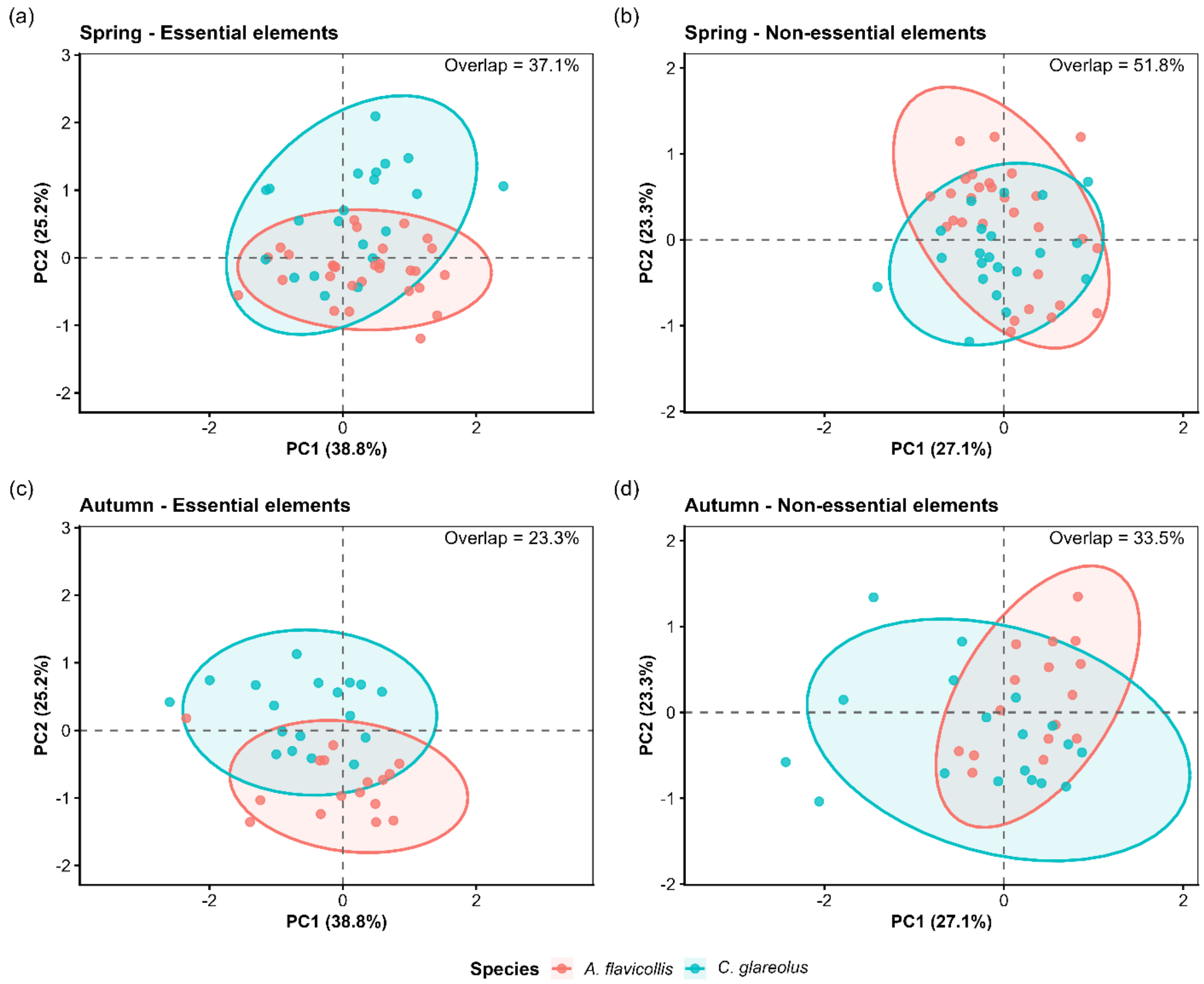


**Figure S7**. Seasonal patterns of interspecific partitioning in essential and non-essential mandibular elementomes. Principal component analyses (PCAs) of *clr*-transformed elemental compositions are shown for essential (a, c) and non-essential elements (b, d) in spring (a, b) and autumn (c, d). Points represent individuals, colours indicate species, and ellipses represent 95% ellipses for each species. Percentages indicate interspecific overlap between species ellipses, calculated as the intersection area relative to their union.

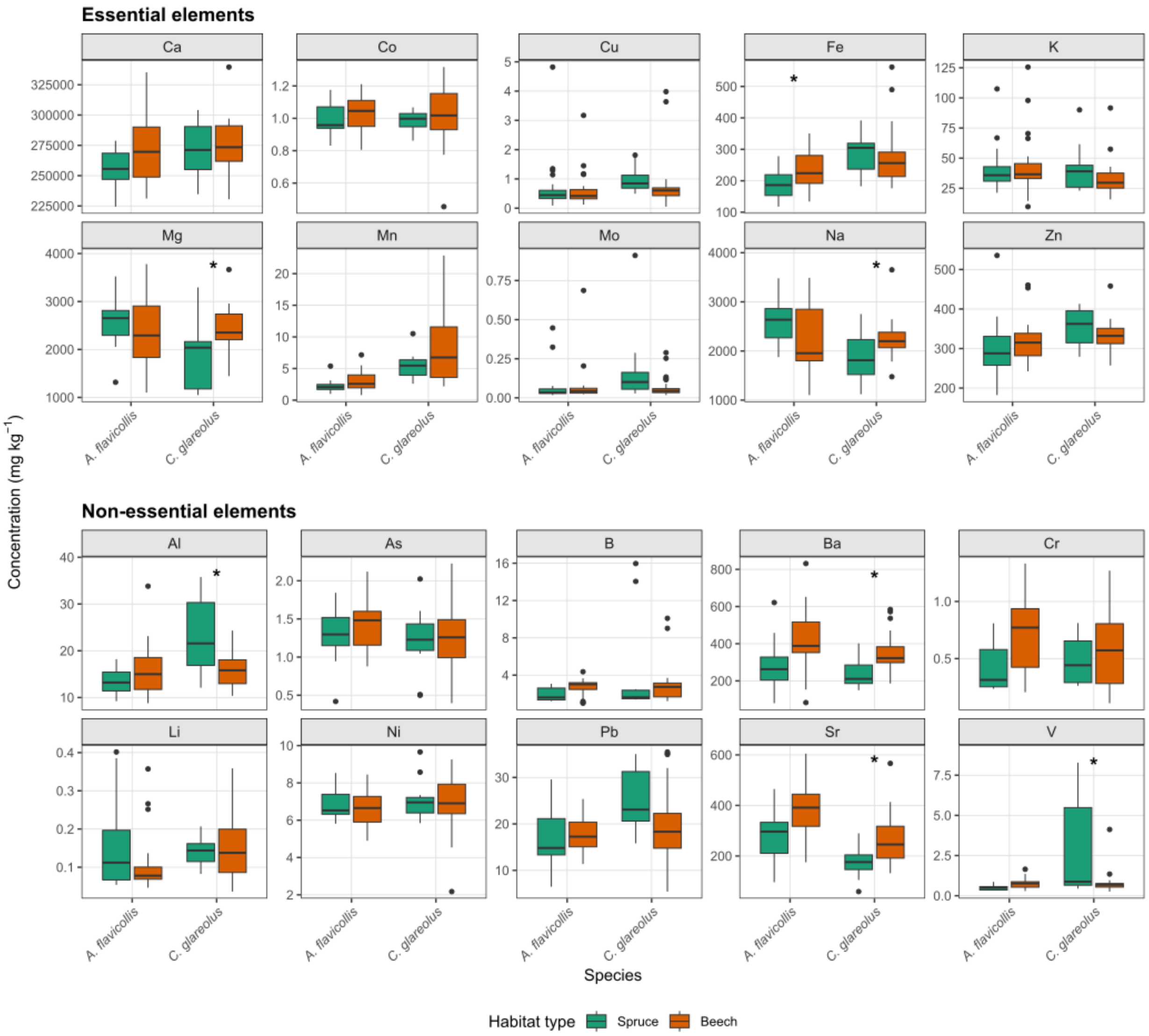


**Figure S8.** Habitat variation in mandibular elemental concentrations of *Apodemus flavicollis* and *Clethrionomys glareolus*. Boxplots show essential and non-essential elemental concentrations by habitat type within each species. Asterisks indicate significant habitat effects from element-specific linear models (Table S11).
